# Simulation of the mechanics of actin assembly during endocytosis in yeast

**DOI:** 10.1101/518423

**Authors:** Masoud Nickaeen, Julien Berro, Thomas D. Pollard, Boris M. Slepchenko

**Affiliations:** Richard D. Berlin Center for Cell Analysis and Modeling, Department of Cell Biology, University of Connecticut Health Center, Farmington, CT 06030, USA; Department of Molecular Cellular and Developmental Biology, University of Connecticut Health Center, Farmington, CT 06030, USA; Departments of Molecular Biophysics, Biochemistry and Cell Biology, University of Connecticut Health Center, Farmington, CT 06030, USA; Nanobiology Institute, Yale University, New Haven, CT 06520, USA

**Keywords:** endocytosis, actin meshwork, mathematical model, yeast data, visco-active fluid, force generation

## Abstract

We formulated a spatially resolved model to estimate forces exerted by a polymerizing actin meshwork on an invagination of the plasma membrane during endocytosis in yeast cells. The model is a continuous approximation tightly constrained by experimental data. Simulations of the model produce forces that can overcome resistance of turgor pressure in yeast cells. Strong forces emerge due to the high density of polymerized actin in the vicinity of the invagination and because of entanglement of the meshwork due to its dendritic structure and crosslinking. The model predicts forces orthogonal to the invagination that would result in a flask shape that diminishes the net force due to turgor pressure. Simulations of the model with either two rings of nucleation promoting factors as in fission yeast or a single ring of nucleation promoting factors as in budding yeast produce enough force to elongate the invagination against the turgor pressure.

## Introduction

Assembly of actin filaments at sites of endocytosis is necessary for invagination of the plasma membrane in both budding and fission yeast (Aghamohammadzadeh and Ayscough, 2009; Basu et al., 2014). The transient accumulation of actin filaments around the invaginating plasma membrane is called an “actin patch.” Patches form in ~10 s, peak and disappear over ~10 s. Polymerizing actin is believed to produce the forces required to form a tubular invagination of the plasma membrane with a clathrin-coated hemisphere at the tip (Kaksonen and Roux, 2018). Force is required to overcome the very high turgor pressure in yeast cells, which is estimated to be on the order of 10 atm in fission yeast (Basu et al., 2014). This amounts to a force on the order of 3,000 pN on a typical endocytic tubule (Carlsson, 2018). Previous modeling studies concluded that actin polymerization alone is unlikely to generate such a force, and various additional mechanisms were proposed (Scher-Zagier and Carlsson, 2016; Lacy et al., 2018).

We used simulations of mathematical models to estimate the forces exerted on an endocytic, plasma membrane tubule by a surrounding network of actin filaments. In our model, mechanics of the filamentous meshwork is coupled to a detailed description of actin nucleation and polymerization (Berro et al., 2010). We assumed that nucleation-promoting factors (NPFs) reside on the membrane tubule and stimulate Arp2/3 complex to nucleate branched actin filaments. Simulations of the model constrained by experimental parameters yielded dense networks of actin filaments around the tubule in the vicinity of the NPFs. Entanglement of the branched filaments makes the network highly viscous, so that the energy released during the polymerization generates forces sufficient to work against the turgor pressure and elongate the nascent invagination.

The elongating invaginations were simulated with either one or two narrow bands of NPFs around the membrane tubule. Fission yeast have two rings of NPFs, one that remains in the initial position at the base of the invagination, while the other moves with the tip of the tubule (Arasada and Pollard, 2011; Arasada et al., 2018). Budding yeast have one ring of NPFs that remains near the base of the invagination.

## Model

### 1. Generalized description of the biochemistry and physics of the expanding actin filament network

The model of the actin filament network is formulated in a continuous approximation, such that the distribution of filaments in the patch is characterized by a continuous density of actin subunits *ρ*(**x**, *t*), which is a function of location **x** and time *t*. The peak number of ~6,500 actin subunits per patch in fission yeast (Sirotkin et al., 2010) suffices for a continuous formulation to provide reasonably accurate results. This large number makes a discrete stochastic approach logistically burdensome, though such an approach would otherwise be appropriate, given submicron sizes of endocytic patches (Mund et al., 2018). Our model indicates that the large number of polymerized subunits confined in submicron volumes results in subdomains of highly concentrated actin filaments in the vicinity of one or two rings of nucleation promoting factors (see *Results*).

We describe filamentous actin as a visco-active fluid (Kruse et al. 2005; Prost et al., 2015). In a viscosity-dominated environment, a balance between active and dissipative forces governs the mechanics of actin filament networks. The active repulsive stress, originating from the impingement of polymerizing subunits on existing filaments, is elastically stored in the meshwork, causing it to expand with velocities limited by dissipation due to viscosity of the meshwork.

Mathematically, a force-balance equation requires that the divergence of the total stress tensor be zero everywhere in the fluid (Kruse et al. 2005): 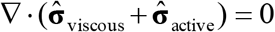. Here, the viscous stress tensor is 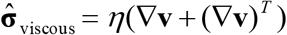, where the viscosity coefficient *η* is a function of the local densities *ρ* and local average length of actin filaments, *L* : *η* = *η*(*ρ, L*) (Doi and Edwards, 1998). Note that because *ρ* varies in space, actin velocities are not subjected in our model to the incompressibility condition. The density of actin subunits, however, has an upper limit due to excluded volume, as explained further in this section.

The active stress tensor is approximated as isotropic: 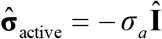, where 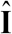 is the unit tensor and *σ_a_*, which is a function of *ρ* (Satcher and Dewey, Jr., 1996; MacKintosh et al., 1995; Gardel et al., 2003), can be interpreted as the energy per unit volume stored in the meshwork during polymerization. Hydrostatic pressure is not included in the force-balance equation in our model, because the mechanics of the actin filament network decouples from mechanics of the cytoplasm. Indeed, the viscous drag exerted on actin filaments by the cytoplasm is much weaker than the intrinsic viscous forces due to direct contacts of the filaments and can thus be ignored (Nickaeen et al., 2017). Technically, the repulsive active stress can be viewed as playing a role of pressure in our model. Overall, the equation governing **v**(**x**, *t*) is written as

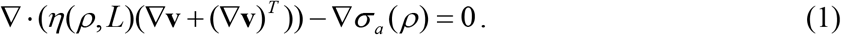

Eq (1) is coupled with the spatiotemporal dynamics of the molecules regulating actin filament assembly. In both types of yeast cells, proteins called nucleation promoting factors (NPFs) initiate the assembly of the actin filament networks by stimulating Arp2/3 complex to nucleate new actin filaments on the sides of existing filaments, forming a dendritic network. The model includes a spatial description of actin nucleation and polymerization that follows a kinetic model used by Berro et al. (Berro et al., 2010). The latter consists of rate equations detailing actin filament nucleation, polymerization and aging, as well as capping the polymerizing (barbed) filament ends and severing of aged filaments by cofilin. Simulations of the model using protein concentrations measured in cells (Berro et al., 2010) adequately describe experimentally measured time courses of the appearance and disappearance of patch proteins (Sirotkin et al., 2010). The rate constants giving good fits of the simulations to the experimental data were larger than expected from biochemical measurements owing to excluded volume effects in cells. Utilizing rate constants and equations of Berro et al. integrates measurements of actin kinetics in our model.

The actin density *ρ* is determined by concentrations of all of the species of actin in an actin patch. These species include newly polymerized ATP-bound subunits (‘FATP’), subunits aged by ATP hydrolysis and phosphate dissociation (‘FADP’), and subunits bound by cofilin (‘FCOF’) as shown in the reaction diagram in Fig. 1. In our model, *ρ* also includes concentrations of the filaments barbed-ends, both active and capped (‘BEa’ and ‘BEc’, respectively), and slowly depolymerizing pointed ends (‘PE’). Overall,

**Figure 1.**
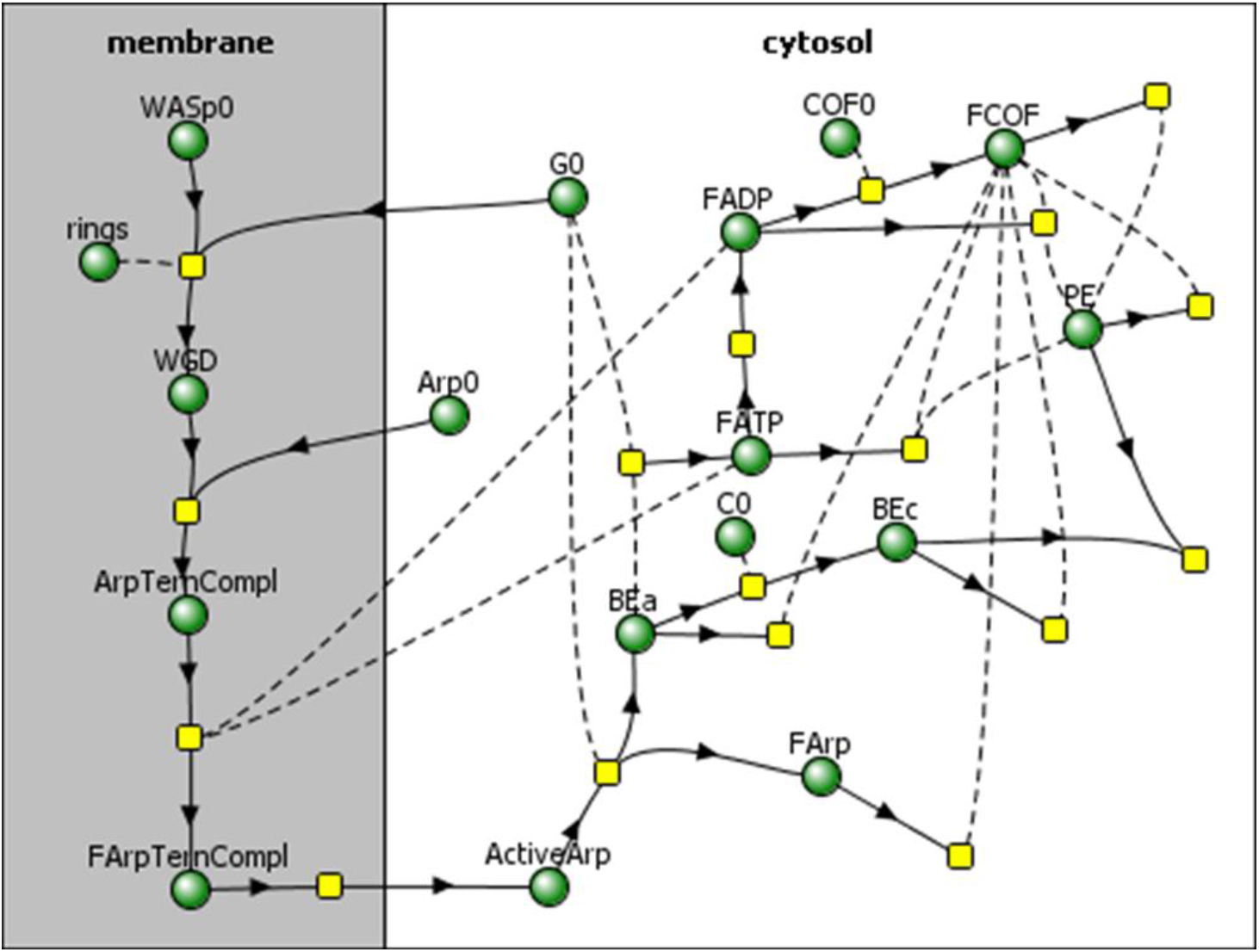
Reaction diagram corresponding to the kinetic model by Berro et al. (Berro et al., 2010), with added partitioning of species between membrane and cytosol. Directions of arrows towards or from reaction nodes (yellow squares) determine roles of species (green circles) in a particular reaction as reactants or products, and reactions without products describe disappearance of reactants from the patch. Species connected to reactions by dashed curves act as ‘catalysts’, i.e. they are not consumed in those reactions.

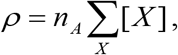

where *x* stands for FATP, FADP, FCOF, BEa, BEc, and PE, and [*X*] is the concentration of molecule *X* in μM; the prefactor *n_A_* converts the concentration in μM into the density expressed in molecules per μm^3^ (*n_A_* = 602 μm^-3^/μM).

All concentrations [*X*], with the exception of [ActiveArp], are governed by reaction-transport equations of the following type,

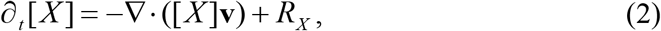

where the first term in the right-hand side describes the flow of *X* with velocity **v** and *R_X_* is the sum of rates of all reactions affecting *X*. The next section describes the equations for [ActiveArp]. Functional forms of *R_X_* and parameters are from (Berro et al., 2010), except the on- and off-rate constant of polymerization, capping, and cofilin binding, which were modified by the factor (1 − *ρ*/*ρ*_max_)^*λ*^ that accounts for the effect of excluded volume. This factor ensures that the abovementioned processes slow down as *ρ* approaches *ρ*_max_ = (4*πδ*^3^/3)^−1^, where *δ* = 2.7 nm is the subunit radius, so that *ρ* never exceeds *ρ*_max_. The functional form used follows from the dependence of molecular diffusivities on the excluded volume (Novak et al., 2011). The parameters used in computations are *λ* = 0.5 (Novak et al., 2009) and *ρ*_max_ = *n_A_* · 19.5 mM (see *Methods* in Supplemental Material).

Reaction steps that lead to formation of ActiveArp occur on the surface of the membrane (Fig. 1) and involve the dimers WASp - G-actin (WGD), Arp2/3 ternary complexes consisting of Arp2/3 bound to WGD(ArpTernCompl), and activated Arp2/3 ternary complexes (FArpTernCompl). These reactions are described by rate equations,

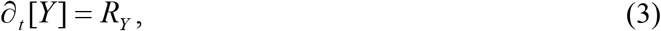

where [*Y*] is the surface density of a membrane-bound protein *Y*. Note that while these variables are governed by ordinary differential equations, they also depend on spatial coordinates, given that *R_Y_* are nonzero only at the locations of NPFs (see below) and *R*_ArpTernCom1_, is dependent on [FATP] and [FADP] near the plasma membrane.

### 2. Coupling the expansion of the actin filament network to the membrane invagination

Eqs (1) and (2) are solved in a sufficiently large neighborhood of the invagination, denoted *Ω* in Fig. 2. The plasma membrane *Γ* includes the invagination. As mentioned above, Eqs (3) are solved on the parts of the invagination occupied by NPFs. Fission yeast assemble two rings containing different NPFs around the invagination of the plasma membrane (red bands in Fig. 2). Both zones start near the cell surface at the neck of the invagination. One ring is stationary, while the other moves with the tip of the invagination, where it is assumed to be attached to a hemisphere of the protein clathrin. Budding yeast has a single zone containing both types of NPFs, which remains at the base of the invagination.

**Figure 2.**
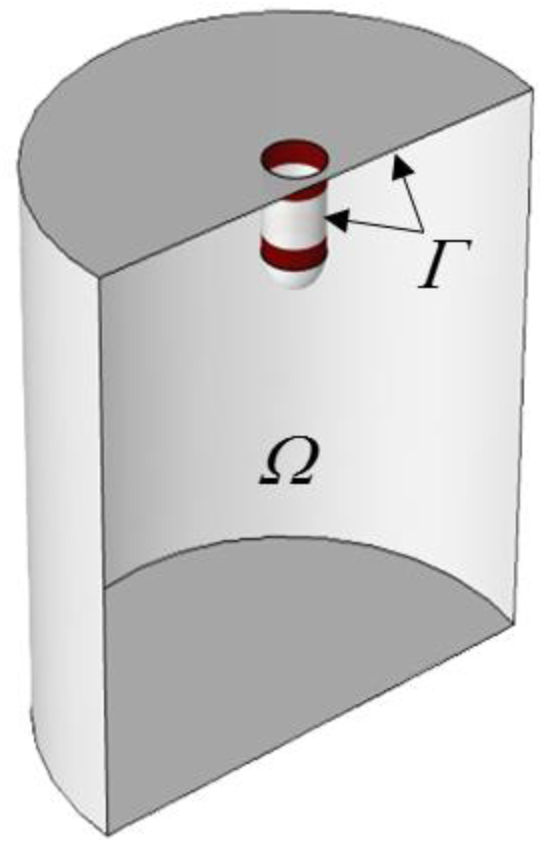
Computational domain, *Ω*, and plasma membrane, *Γ*, including invagination. Two rings of nucleation-promoting factors are shown in red. When the invagination elongates, both *Γ* and *Ω* change with time.

We assume that the actin patch assembly is preceded by formation of an initial invagination. We do not model this process, which involves formation of a coated pit of plasma membrane associated with clathrin molecules and adapter proteins (Arasada and Pollard, 2011; Chen and Pollard, 2013). For this study, we assume the existence of the initial invagination with a depth sufficient to accommodate two adjacent rings of NPFs. The next section describes the shape and size of the initial invagination used in simulations.

Actin filaments polymerizing around the initial invagination are constrained by the plasma membrane, which is pressed against the stiff cell wall. This causes the actin filament network to expand inward from and laterally along the cell surface. The expanding meshwork exerts a drag on an initial invagination and moves the invagination further inward. It is believed that the drag occurs because of binding of actin filaments to the protein coat of the invagination (Lacy et al., 2018), though little is known about the biochemistry of this binding. The connection between the actin meshwork and the plasma membrane is included in the model as a condition that the membrane and the adjacent actin filaments move with the same velocities: (**v** − **u**)|_*Γ*_ = 0, where **u**|_*Γ*_ are the velocities of the points of the membrane. This condition is consistent with the treatment of viscous fluids at interfaces with adjacent media in continuum mechanics (Landau and Lifshitz, 1987). Mathematically, it serves as a boundary condition for Eq (1) at *Γ*. The conditions at other boundaries of the computational domain were zero-stress, though they did not affect the solution significantly, since *Ω* was substantially larger than the size of the invagination (see *Methods* in Supplemental Material).

The net force exerted on the endocytic invagination is obtained by evaluating an integral of the tangential force density, 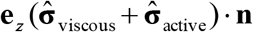 over the surface of the invagination *S*:

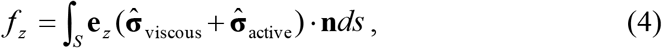

where **n** is the outward normal vector to *Γ* (directed from *Γ* towards the interior of *Ω*), **e**_z_ is the unit vector orthogonal to the cell wall and *ds* is the infenitesimal surface element (Landau and Lifshitz, 1987). The *Results* section considers in detail the rheological data for actin networks that are critically important for the constitutive dependences *σ_a_* = *σ_a_* (*ρ*) and *η* = *η*(*ρ, L*) used in Eq (1).

Eq (2) is subject to zero-flux boundary conditions at *Γ* for all *X*, except for ActiveArp, for which there is an incoming flux from the rings that describes the detachment from the membrane of active filament-bound ternary complex (‘FArpTemComp?), see Fig. 1. The magnitude of the corresponding flux density is equal to the detachment rate *R*_FArpTernComp1->ActiveArp_ |_*γ*_rings__, where *γ*_rings_ denotes the zones of *Γ* occupied by the rings (see Fig. 2 and *Methods* in Supplemental Material). The existence of a nonzero influx of ActiveArp requires modification of the transport term in Eq (2) for this variable. Indeed, given the boundary condition for **v**, pure advection is generally incompatible with a nonzero influx, resulting in unphysical Dirac-delta singularities. The inconsistency is resolved by taking into account that the detachment of the ternary complex from the membrane inherently involves diffusion. Adding the diffusive term restricted to the vicinity of the rings, we arrive at:

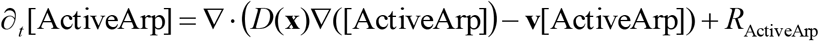

and a corresponding boundary condition, (*D*(**x**)Δ([ActiveArp]) + *R*_FArpTernComp1->ActiveArp_)|_*γ*_rings__ = 0, where *D*(**x**) is nonzero only in the vicinity of the rings (see *Methods* in Supplemental Material).

At all the other boundaries of the computational domain, Eq (2) was subject to the outflow boundary conditions. As we have noted in the context of Eq (1), the type of these boundary conditions does not really matter, because so long as the size of *Ω* is sufficiently large, they do not affect the solution in any significant way (see *Methods* in Supplemental Material).

### 3. Simulations of the models

Eqs (1-3) coupled with respective boundary conditions were solved numerically. Importantly, when the membrane elongates, *Γ* and *Ω* in Fig. 2 are changing: *Γ* increases and *Ω* decreases, so the model must be solved in a domain with a moving boundary (see *Methods* in Supplemental Material). Note that the concentrations of molecules with names followed by zero in Fig. 1 are constants and the surface density of the nucleation-promoting factors, WASp0, is uniform within the rings and varies over time as a bell-shape curve (Sirotkin et al., 2010; Berro et al. 2010). The initial values of all other concentrations and **v**(**x**,0) were set to zero, except for [FADP], [BEa], and [PE], which were assigned small initial values, corresponding to a small number of seed filaments (Chen and Pollard, 2013).

The geometry of the initial invagination was a cylinder with radius 30 nm capped with a hemisphere of the same radius. The initial length of the cylindrical part was 40 nm, accommodating two 20-nm wide rings positioned next to each other. It was assumed, for simplicity, that during elongation, the invagination preserves its (sphero)cylindrical shape and is infinitely rigid i.e. that all points of the tubular membrane have the same instantaneous velocities collinear with the axis of the cylinder. Realistically, the invaginations are not infinitely rigid. Indeed, electron micrographs showed the endocytic invaginations of budding yeast are of flask shape (Kukulski et al., 2012). Our model yields forces orthogonal to the tubule distributed in a way that would produce such a shape (see Fig. 6).

We computed the time-dependent magnitude of these velocities assuming a linear force-velocity relationship (Peskin et al., 1993),

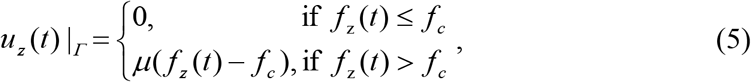

where *f_z_*(*t*) is the force exerted on the invagination at time *t*, defined by Eq (4), *f_c_* is the critical force due to turgor pressure, and *μ* is a given mobility coefficient (see *Methods* in Supplemental Material and *Results*).

## Results

### 1. Parameterization of Eq (1) and salient properties of the model

We begin with a description of constitutive relations for active stress and viscosity of actin meshwork in the absence of branching and crosslinking. Measurements of the viscoelasticity of filaments of purified actin can explain how the active stress and viscosity of the meshwork depend on its density and the properties of the filaments. Rheological data usually include information about dynamic (i.e. frequency-dependent) ‘storage’ and ‘loss’ moduli, denoted as *G′*(*ω*) and *G″* (*ω*), respectively (Wirtz, 2009). The active stress, *σ*_a_, which is determined by the energy released during polymerization and elastically stored in the meshwork, should be proportional to *G*′. For overlapping actin filaments, *G′*(*ω*) scales with actin density *ρ* as ∝ *ρ*^2^ for any *ω* (Gardel et al., 2003). We therefore assume *σ*_a_ = *κ*_active_*ρ*^2^, where the proportionality coefficient *κ*_active_ depends on the extent of branching and crosslinking.

Obtaining a constitutive relation for viscosity *η* is not as straightforward. Based on polymer physics, it is expected to be of the form, *η ∝ ρ^α^L^β^*, where *L* is the polymer length and exponents *a* and *β* depend on whether the polymer is flexible or rigid and whether the solution is dilute or concentrated (Doi and Edwards, 1998). For concentrated solutions of certain flexible chemical polymers, measurements yielded *α* = 4-5 and *β* ≈ 3.5, in agreement with theoretical results. Note that the same theory predicts that the viscosity of a polymer solution is always proportional to viscosity of a solvent; this is based on the assumption that the cross-sectional area of a polymer is vanishingly small. While this assumption is adequate for chemical polymers, it does not apply to a biopolymer meshwork, where the viscosity originates from direct interactions between filaments and is essentially independent of viscosity of the medium. It is intuitive to assume that viscosity of overlapping actin filaments increases as a function of the number of contacts made by the filaments and how long these contacts ‘slide’ along the filaments. The average number of contacts a given filament makes with its neighbors can be estimated as the average number of subunits per volume occupied by a filament, i.e. ~ *ρNδ*^3^, where *N* is the average number of subunits per filament and *δ* is the radius of the actin subunit, as defined earlier. The contact density is then obtained as a product of the number of contacts per filament and the number of filaments per unit volume. The latter is *ρ*/*N*, so that the density of contacts is ~ *ρ*^2^*δ*^3^. Assuming further that for the rod-like filaments, the ‘lifetime’ of a contact is proportional to the number of subunits in a filament *N*, we arrive at *η* ~ *ρ*^2^*δ*^3^*N = ρ*^2^*δ*^2^*L*, or

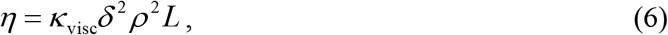

where the proportionality coefficient *κ*_visc_ can depend on the structural properties of an actin meshwork, such as branching or crosslinking.

We corroborated the constitutive relation of Eq (6) by estimating *η* from rheological data for filaments of purified actin. The estimation of *η* is complicated by the fact that solutions of actin filaments are non-Newtonian fluids with viscosities depending on the shear rates (Buxbaum et al., 1987). This was approximated by deriving *κ*_visc_, treated as a constant, from *G′*(*ω*) and *G″*(*ω*), with *ω* close to the shear rates in actin patches, which are ~ 1 Hz (see the end of this subsection for more details). It is also important to note that the shear viscosity of the meshwork differs from *η*′(*ω*) = *G′*(*ω*)/*ω* (Cox and Merz, 1958; Wirtz, 2009). The effective shear viscosity is often well approximated by an empirical Cox-Merz rule *η* = ω^−1^(*G′*^2^(*ω*) + *G′*^2^(*ω*))^12^, with *ω* being identified with the shear rate (Cox and Merz, 1958). In what follows, values of *η* were computed by applying the Cox-Merz formula to the moduli measured at *ω* = 1 Hz.

The length dependence in Eq (6) is close to *η* ∝ *L*^0.7^ as proposed by Zaner and Stossel (Zaner and Stossel, 1983), who measured dynamic moduli of solutions of overlapping actin filaments with controlled lengths and applied the Cox-Merz rule to compute *η*. More recent data by Kasza et al. points to a linear dependence, *η ∝ L* (Kasza et al., 2010). These authors measured *G′* (*ω*) and *G*″(*ω*) of overlapping actin filament networks prepared with a fixed actin concentration and varying filament lengths and concentrations of linkers. Extrapolation of the data of (Kasza et al., 2010) to a zero cross-linker concentration gives the filament length dependence of *η* without crosslinking. Specifically, the data points of Figure 4c in (Kasza et al., 2010), corresponding to *ω* = 1 Hz, were extrapolated to the linker-to-actin concentration ratio *R* = 0 by approximating the increase in viscosity due to cross-linking as ∝ *(RL*) (McFadden et al., 2017). Fig. 3, which includes data for*R =* 0 of Figure 4a in (Kasza et al., 2010), shows the dependence of *η* on filament length in the absence of crosslinking or branching.

**Figure 3.**
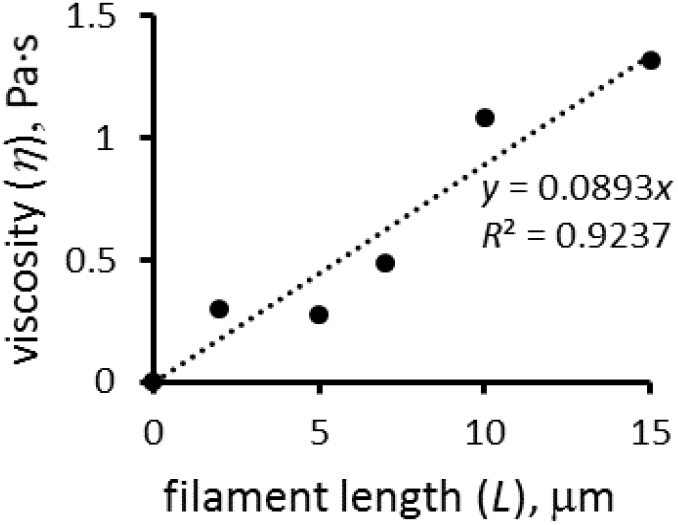
Viscosity of actin filament meshwork as a function of mean filament length at *ρ/n_A_* = 12 μM. Extrapolated from data of (Kasza et al., 2010).

**Figure 4:**
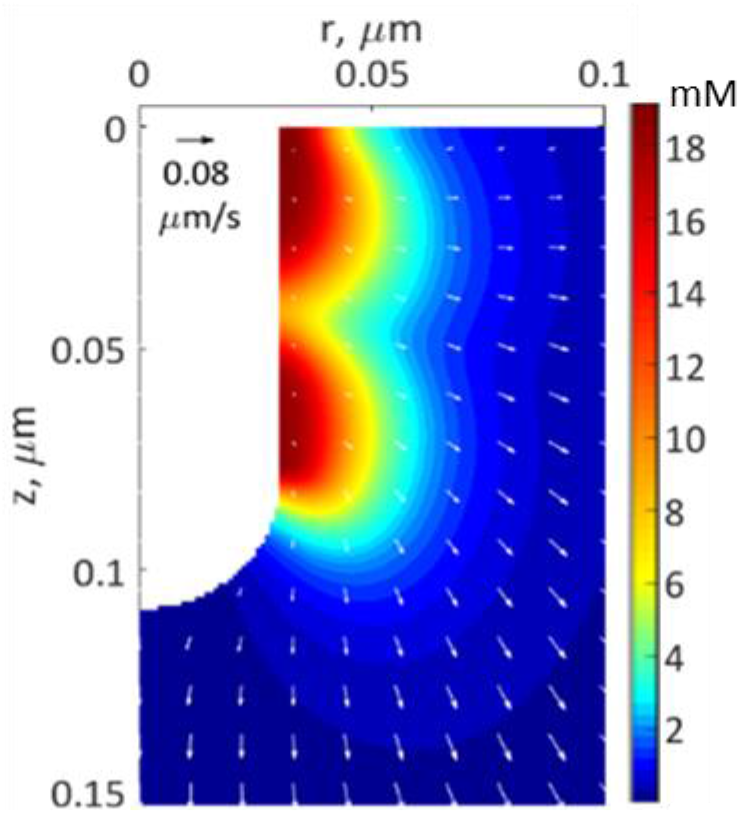
A snapshot from a simulation of an elongating endocytic invagination shown for *r - z* cross-section of 3D geometry. The extracellular space is white. The false color shows the density distribution of actin filaments, and the arrows show the local velocities of their movements at the peak of actin assembly. See Fig. 7C for snapshots at other time points.

To confirm the quadratic *ρ* dependence of Eq (6), one would need rheological data for actin filament samples with a fixed filament length and a range of actin concentrations. The data closest to these requirements are for *G′*(*ω*) and *G*″(*ω*) of pure actin filaments without branching or crosslinking at concentrations of 1 mg/mL and 0.3 mg/mL (Gardel et al., 2003).

Measurements at *ω* = 1 Hz yielded *η ∝ ρ^α^* with *α* = 1.98. Eq (6) also yields plausible average filament lengths, 15 μm and 12 μm, based on the data for pure actin filaments reported in (Sato et al., 1987) and (Mullins et al., 1998), respectively. These values were obtained using *κ*_visc_ for pure actin filaments that was estimated by applying Eq (6) to data points in Figure 4a of (Kasza et al., 2010) corresponding to ***R*** = 0 (open and filled triangles) and *ω* = 1 Hz. In this experiment, *L* = 15 μm, [*F*_tot_] = *ρ*/*n_A_* = 0.5 mg/mL=12 μM, and the respective viscosity *η*, computed by the Cox-Merz rule, is 1.32 Pa·s, yielding *κ*_visc_*n_A_* ≈ 0.14 Pa·s/μM.

Note that Eq (6) holds only for overlapping filaments, i.e. for dense actin networks of sufficiently long filaments, such that (*ρN*^2^)^1/3^ δ > 1 (Doi and Edwards, 1998). This condition is most certainly violated at early stages of patch assembly, when only few short filaments are present. In this limit, *η* is expected to be a multiple of solvent viscosity and ∝ *ρ*. Because noticeable stresses and shear rates are generated only after filaments begin to overlap, the two regimes were bridged in our computations by using a simple ‘interpolation’ formula, that crosses over to Eq (6) when the condition for the filament overlapping is met,

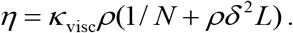

In this formula, the number of subunits per filament *N* was computed as [*F*_tot_]/([BEa] + [BEc]), where [BEa] + [BEc] is equivalent to local filament number density, and the filament length is *L = Nδ*, as above.

Substituting the constitutive relations *σ _a_* (*ρ*) = *κ*_active_*ρ*^2^ and *η*(*ρ, L*) = *κ*_visc_*ρ*^2^δ^2^ *L* in Eq (1) yields

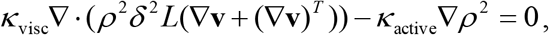

from which it follows that both actin densities *ρ*(**x**, *t*) and velocities **v**(**x**, *t*) are controlled by the ratio *κ*_active_ *κ*_visc_, rather than separately by *κ*_actlve_ and *κ_visc_* (as defined earlier, here and below vectors x denote spatial coordinates of a location in the cell). We confirmed, by solving the model numerically with varying *κ*_active_ and *κ*_visc_, that **v**(**x**, *t*) did not change beyond numerical error, when both coefficients were varied proportionally. Also in agreement with the prediction, we found that *κ*_active_ *κ*_visc_ controls a maximum number of polymerized subunits in a patch 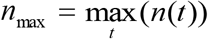, where 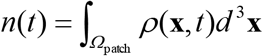 is the number of subunits at time *t* in the *t* patch volume *Ω*_patch_ occupied by the invagination and surrounding network of actin filaments. Modeling an elongating cylindrical invagination with varying *κ*_active_ *κ*_visc_ (see *Dynamics of the invagination during elongation*), we found that the ratios 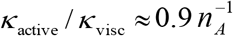 Hz/mM result in *n*_max_ close to the experimental numbers. For example, the ratio 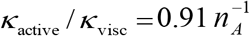 Hz/mM yields the maximum number of 6450 subunits inside a cylinder *Ω*_patch_ of radius 0.16 μm and length 0.32 μm enveloping the endocytic tubule. The ratio *κ*_active_ *κ*_visc_ constrained by the experimental *n*_max_, in turn, determines actin velocities *v*(**x**, *t*) and the corresponding shear rates, which are found to be ~ 1 Hz (see below).

Fig. 4 depicts a snapshot of a solution of the model with 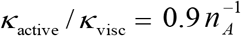 Hz/mM showing distributions of actin density (pseudo-colors) and actin velocities (white arrows) for an *r - z* section (*r* and z are cylindrical coordinates) at a time when the rings on an elongating invagination have separated. The solution yields two zones of actin filaments, which are particularly dense in the vicinity of the rings. Note that even though the two rings were identical in size and density of nucleation-promoting factors, the actin filament density is higher near the plasma membrane, owing to the inhomogeneity of active barbed ends whose transport is restricted by the rigid cell wall surrounding the plasma membrane. The gradient of actin density then results, as expected, in a net tangential force directed towards the tip of the invagination. Note that radial and tangential components of actin velocities in the vicinity of the invagination are ~0.02 μm/s, yielding patch diameters of ~100-200 nm, consistent with experimental data (Berro et al., 2010; Arasada et al., 2018). The solution also indicates (data not shown) that tangential components of actin velocity vary significantly in the normal direction over distances ~0.02 μm from the membrane, yielding shear rates of ~1 Hz, as mentioned above.

Control of the shear rates and actin densities by *κ*_active_ / *κ*_visc_ has another consequence: for a given *n*_max_, the force exerted on the invagination depends on *κ*_visc_ (or alternatively on *κ*_active_, given that *κ*_active_ / *κ*_visc_ is fixed). Mathematically, this is seen upon substitution of the constitutive relations in Eq (4). Qualitatively, the tangential force exerted on the invagination, which largely originates from the viscous stress, is locally defined by a product of viscosity and shear rates. Since the latter are fixed by the known *n*_max_, this leaves the tangential force to be directly proportional to *κ*_visc_. We confirmed this assertion computationally by solving the model with constant *κ*_active_ / *κ*_visc_ over a range of *κ*_visc_.

### 2. Patch assembly can generate pushing forces comparable to turgor pressure in fission yeast

We use the model with 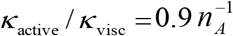 Hz/mM to determine the *κ*_visc_ required to generate forces sufficient to exceed the turgor pressure. For this, we solved the model in a fixed geometry with a shape of the initial invagination with a radius of 30 nm. Supplemental Figure 1 shows the density distributions and velocities of the actin filament network. A tangential force of ~2500 pN is sufficient for such an invagination to withstand turgor pressure of about 8.75 atm after ~10 s of patch assembly. This force requires 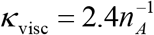 Pa·s/μM, which is ~17-fold larger than 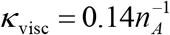 Pa·s/μM of actin filaments alone.

Two factors in patches contribute to a higher viscosity than actin filaments alone. First, the meshwork is highly entangled due to the high density of branching. For example, the viscosity of 24 μM of actin filaments at a shear rate of *ω* = 1 Hz was more than 7-fold higher when polymerized with 0.12 μM of Arp2/3 complex according to Figure 3 in (Tseng and Wirtz, 2004). The molar ratio of Arp2/3 complex-to-actin in these experiments, ***R***_Arp2/3_ = 0.005, was significantly lower than the range of 0.035 and 0.06 observed in actin patches (Berro et al., 2010). Such high ratios ***R***_Arp2/3_ increase the viscosity by at least a factor of 2.5 according to rheological measurements of actin filaments with a range of concentrations of Arp2/3 complex (Mullins et al., 1998). Even without crosslinking the 15-fold higher viscosity results in 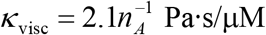 and a pushing force of 2166 pN, enough to withstand 7.7 atm of turgor pressure. Second, actin patches accumulate a very high concentration of the crosslinking protein fimbrin (Berro and Pollard, 2014), which increases the viscosity. Rheological data indicate that the viscosity of actin networks cross-linked by soft (muscle alpha-actinin, filamin) and rigid (avidin-biotin) linkers ranges from few fold to an order of magnitude higher than actin filaments alone without Arp2/3 complex (Wachsstock et al. 1994, Kasza et al. 2010). The properties of actin filaments cross-linked by fimbrin are likely to be in the same range.

Our simulations of patch formation and force generation must satisfy several constraints. For a fixed *κ*_active_ / *κ*_visc_, the increase of *κ*_visc_ implies a similar increase of *κ*_active_ and hence the corresponding increase of *σ*_a_. The latter is limited by free energy released during a polymerization step *k_B_T* ln(*G*_0_ / *G*_crit_), where *G*_0_ is the concentration of actin monomers and the critical concentration 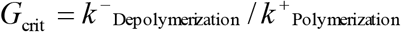 (Footer et al., 2007). For the parameter values used in our model, the upper bound for the stored energy is 6.9 *k_B_T* and the corresponding stalling force is estimated to be, *f*_stall_ = 6.9 *k_B_T/ δ* ≈ 10.5 pN, consistent with published estimates of *f*_stall_ (Lacy et al., 2018). Based on the simulated actin densities *ρ*(**x**, *t*), the maximum energy stored in the patch per subunit is 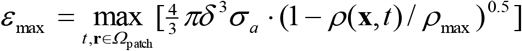, where *σ^a^ = κ*^active^*ρ*^2^(**x**,*t*) (see the previous subsection), and the corresponding maximum force generated by a polymerizing subunit is 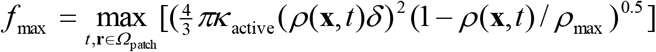 Applying these formulae to the abovementioned solution in the fixed geometry, we find that it satisfies the constraints: *ε*_max_ ≈ 4.9*k_B_T* < 6.9 *k_B_T* and *f*_max_ = 7.5 pN < *f*_stall_.

The ability of a filament to sustain generated forces is another constraint on the system; the force per filament should not exceed the buckling threshold, f_crit_ = *π^2^ EI* /(2L)^2^ (Broedersz and MacKintosh, 2014). In this formula, *E =* 1 GPa is Young’s modulus of the actin filament *I = πa*^4^/4 is the rotational inertia of the filament, where *a* =3.5 nm is the radius of the filament cross-section; and *L* is the filament length. To satisfy the constraint, the force per filament in the vicinity of the invagination must be less than the critical load f_crit_. Using the solution of the model, we first determine the number of filaments in the vicinity of the invagination by integrating the density of barbed ends *ρ*_BE_(**r**, *t*) = *n_A_*([BEa] + [BEc]) in a shell with thickness equal to the shortest filament length (recall that filament lengths are calculated as *L = Nδ*, where *N* = *ρ*(**r**, *t*)/*ρ*_BE_(**r**, *t*)). For the solution with the fixed geometry described above, at the time of the peak of actin assembly, the filament lengths in the vicinity of the endocytic tubule varied from 42 nm to 141 nm, which is consistent with previous estimates (Berro et al. 2010). Integrating *ρ*_BE_(**r**, *t*) in the shell with thickness 42 nm yields 106 filaments, so for this solution the average force per filament is 2541 pN/106 ≈ 24 pN. For roughly half of the filaments having lengths under 105 nm, the critical load is greater than 26 pN. Thus, as expected, the shorter filaments endure the generated force on their own. The longer filaments sustain their share of the load through crosslinking by fimbrin: because the critical load for a bundle of filaments grows roughly as square of the number of bundles filaments, the buckling threshold for a bundle of just two filaments will be at least 100 pN.

We thus conclude that the forces generated during patch assembly can exceed the opposing forces from turgor pressure in fission yeast.

### 3. Dynamics of the invagination during elongation

Once the force exerted on the invagination exceeds the turgor pressure threshold, the invagination will grow inward. The rate of the growth in our model is given by Eq (5): *u_z_*(*t*) = *μ*(*f_z_*(*t*) − *f*_c_). It may seem that the length the invagination can attain during patch assembly is controlled by the mobility coefficient *μ*. However, solving the model in a dynamic geometry with varying *μ* indicates that the final length of the endocytic tubule is virtually insensitive to *μ*. This is true, because the increase of *μ* is mitigated by the drop in *f*_2_ that depends on the shear rates *∂_r_**ν**_z_*, so that the elongation rate *u_z_* does not change appreciably (in computations, we used *μ* = 0.4 nm · s^-^/pN).

The kinetic parameters of actin nucleation and polymerization govern the duration of patch assembly, so the time during which the patch elongates depends on how quickly *f_z_* overcomes the critical threshold *f_c_* from turgor pressure (Fig. 5A). The time before *f_z_* exceeds *f_c_* is shorter as *κ*_visc_ increases, but *κ*_visc_ has an upper bound. The reason is that *κ*_active_ must increase in proportion to *κ*_visc_ while the ratio *κ*_active_ / *κ*_visc_ is limited by a maximum number of subunits in a patch, and *κ*_active_ is limited by the energy constraints considered in the previous subsection.

**Figure 5:**
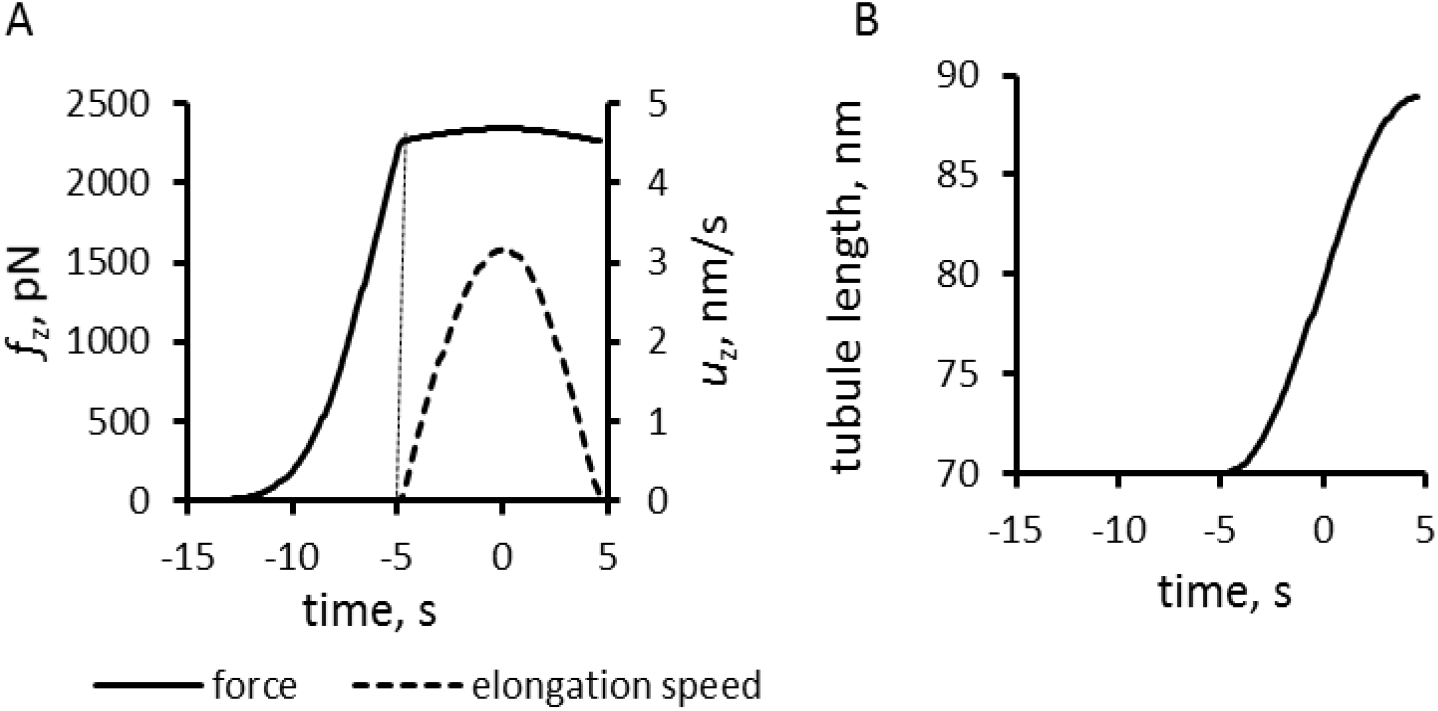
Simulation of the elongation of an endocytic tubule with a fixed threshold corresponding to the turgor pressure of 8 atm. Time zero is the peak of actin assembly. (A) Time course of net tangential force (solid line) and the speed of elongation (dashed line). (B) Tubule length over time.

Solving the model in a geometry allowing the invagination to lengthen freely yields a growing endocytic tubule (Supplemental Movie 1). Fig. 5 illustrates how *f_z_*(*t*), *u_z_*(*t*), and invagination length depend on time with *f*_c_ = 2262 pN, corresponding to a turgor pressure of 8 atm, and with the largest possible 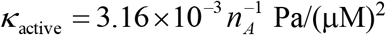 and 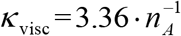 Pa·s/μM. Note that the rate of increase of *f_z_* drops sharply when the exerted force crosses the turgor-pressure threshold (Fig. 5A). Above this threshold, the surface area increases, but *f_z_* plateaus below the values reached in fixed geometry with the same *κ*_active_ and *κ*_visc_, due to the drop of shear rates when the invagination starts to move.

The model produces longer invaginations when taking into account the effects of the forces produced by actin polymerization on the shape of the plasma membrane invagination. The distribution of force density 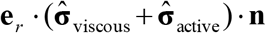 orthogonal to an invagination, shown in Fig. 6A, for the fixed-geometry solution of the previous subsection, suggests that the forces normal to the membrane squeeze the invagination near the plasma membrane and stretch the middle of the invagination. If the tubule is not infinitely rigid, these forces will tend to distort the invagination into flask or ‘head-and-neck’ shape, Fig. 6B, as observed in electron micrographs of budding yeast actin patches (Kukulski et al., 2012).

**Figure 6:**
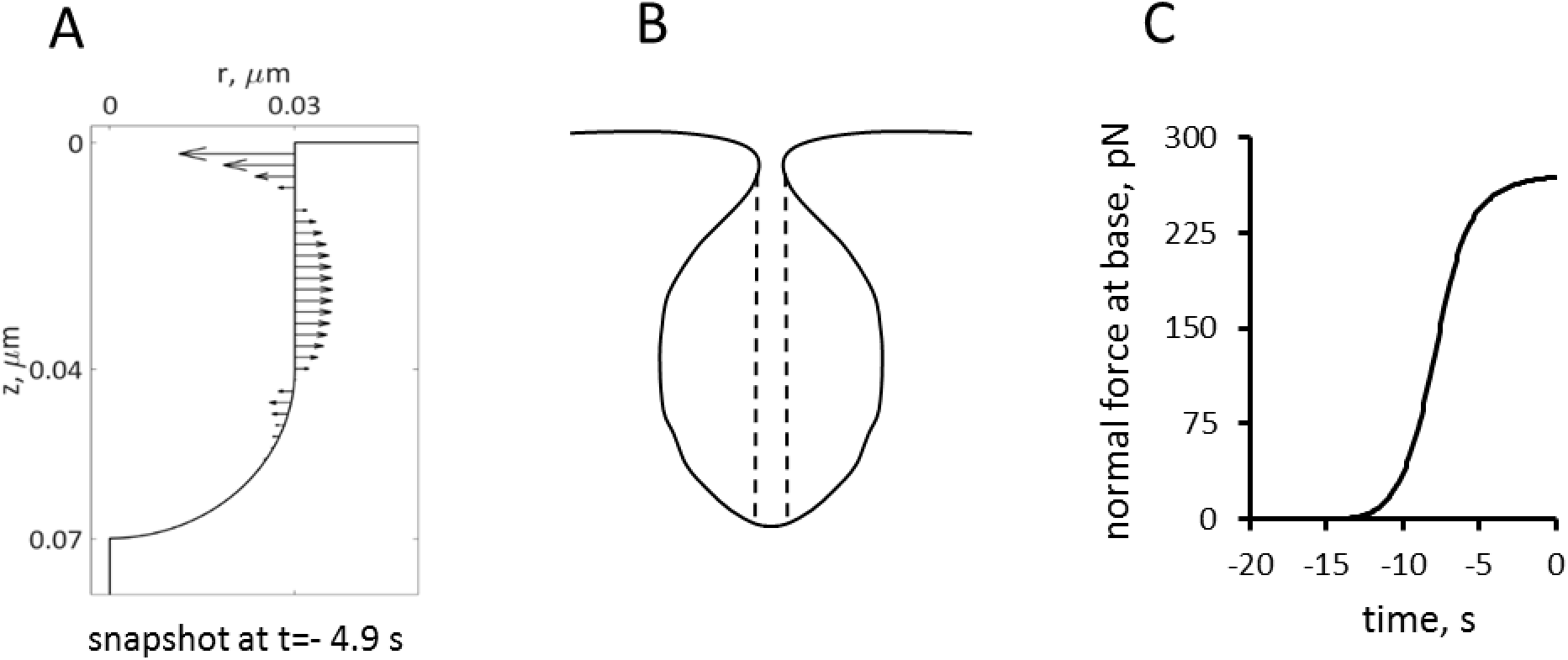
Simulations of the forces exerted by actin assembly normal to the endocytic tubule. (A) Distribution of forces at ≈ 5 s before peak on a tubule of fixed geometry. (B) Rough sketch of a plausible shape if the membrane lining the invagination is flexible. The vertical dashed lines show the area of the pore that determines the force produced by the turgor pressure. (C) Time course of the force normal to the tubule at its base. Time zero is the peak of actin assembly.

Because turgor pressure is isotropic, the net resistance force *f_c_* it produces for the flask shape is proportional to the cross-sectional area of the opening of the invagination delineated in Fig. 6B by dashed lines. This resistance force decreases over time as the normal force from the expanding actin network (Fig. 6B) squeezes the neck of the invagination near the cell surface.

The model with the time-dependent threshold approximated as 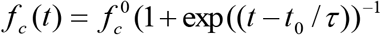, where 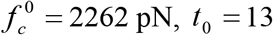 s and *τ* = 1 s, and other parameters as above, yielded a longer invagination than with a rigid tubule (Fig. 7D and Supplemental Movie 2).

**Figure 7:**
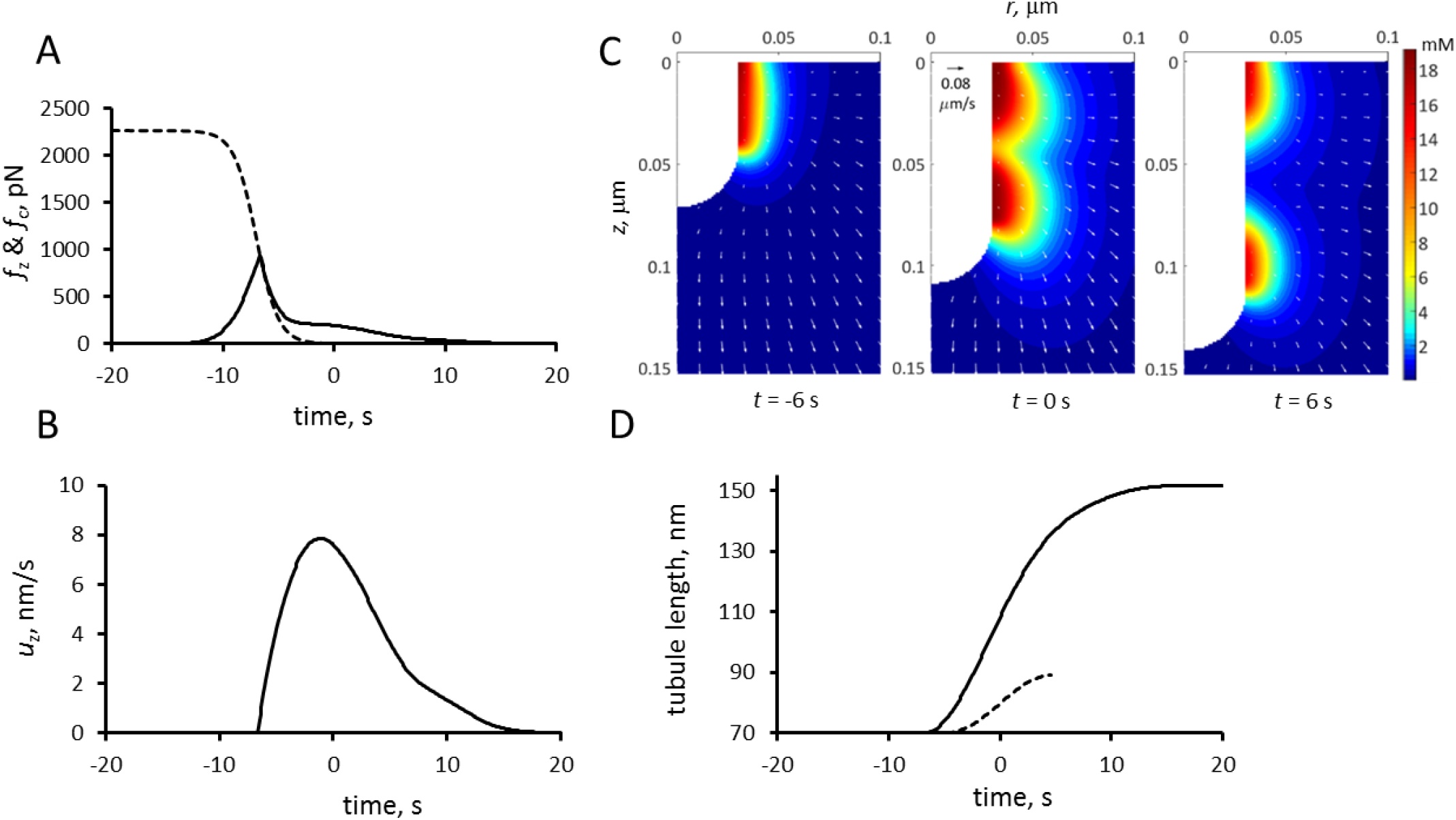
Simulation of endocytic tubule elongation with the force threshold from turgor pressure decreasing with time. Time zero is the peak of actin assembly. (A) Time course of the assumed decrease in force threshold due to turgor pressure, *f*_c_ (dashed curve) and the simulated pushing force, *f_z_* (solid line). (B) Time course of the variation in the speed of invagination, which begins when *f_z_* is greater than *f*_c_. (C) Snapshots of *r-z* sections of the actin filament density around the endocytic tubule and its velocities (arrows); also see Supplemental Movie 2. (D) Comparison of the time courses of tubule elongation with decreasing force from turgor pressure (solid line) with that with a fixed threshold due to turgor pressure in Fig. 5B (dashed curve).

The lengths of modeled invaginations are similar to the distances that actin patch proteins moved from the cell surface in super-resolution movies, taking into account the size of the protein coat around the membrane (Arasada et al., 2018). To illustrate the qualitative agreement between the model and experiment, the simulation data were processed using the protocol adopted by Arasada and Pollard (see *Methods* in Supplemental Material for details), so that the results shown in Fig. 8 can be directly compared with the experimental data (compare Figure 3A-B in (Arasada et al. 2018).

**Figure 8:**
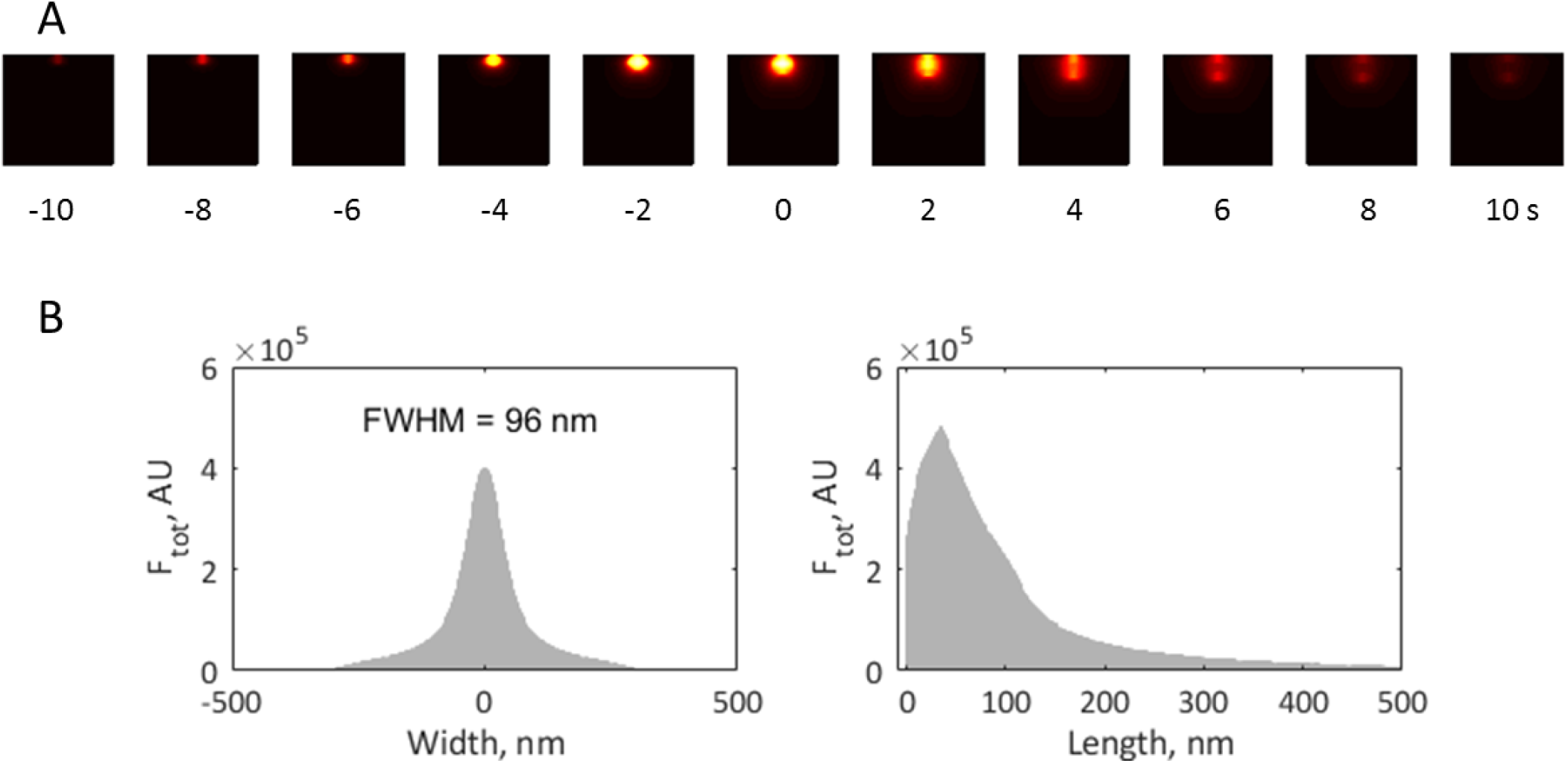
Simulation of elongating tubule with time-dependent force threshold is consistent with experimental data. (A) Heat maps of simulated actin density (see Fig. 7 above) projected on plane of image and subjected to median filtering to mimic 35-nm resolution limit due to convolution with point-spread function, are shown for selected time points. See *Methods* for details of how simulation results were processed for this Figure; see Supplemental Fig. 2 for results before filtering. (B) Width and length distributions of actin density, obtained by integrating results of panel (A) over time, are consistent with experimental data of Arasada and Pollard (Arasada et al., 2018).

We compared the solution of the two-ring model with a fixed threshold *f_c_* with the corresponding solutions of the models, in which all of the NPFs remained at the base or moved together with the tip of the tubule (Fig. 9). For all three versions of the model we used invaginations with the same widths and total numbers of nucleation-promoting factors and ran the simulations with the same initial conditions. The model with the NPFs remaining at the base slightly over-performs the two-ring model. In contrast, the model with the NPFs moving together with the tip generates significantly weaker forces, resulting in a slower movement and much shorter invagination than the two-ring model. These results highlight the importance of the cell wall in supporting the actin meshwork to generate traction forces. The partial absence of such support in the two-ring model is mitigated almost entirely by the repulsion of the two zones of polymerizing actin.

**Figure 9:**
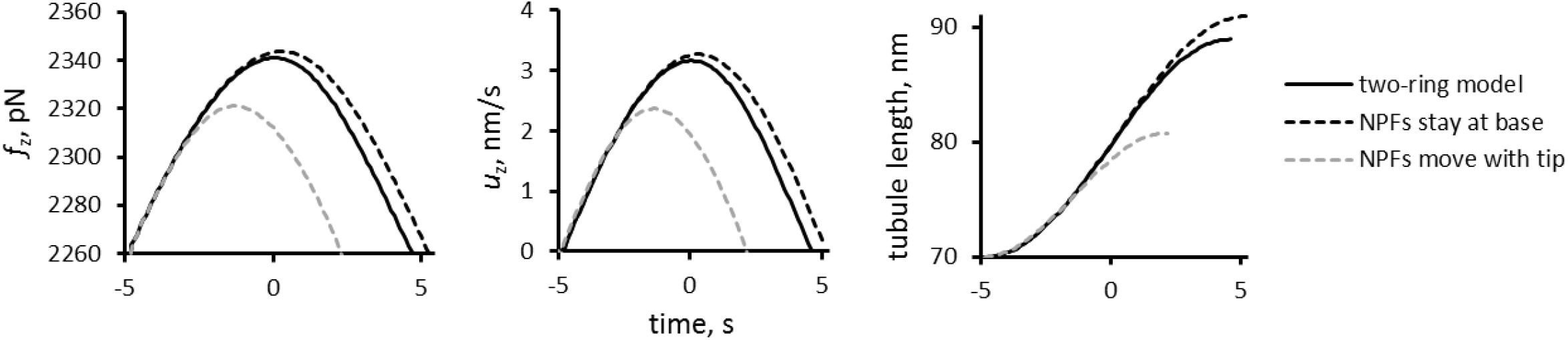
Comparison of the results of simulations of models with three different locations of nucleation promoting factors: solid lines, two-ring model with NPFs at the base and tip of the invagination; dashed line, one ring model where all NPFs stay at base of invagination; and grey dashed line, one ring model with all NFPs at the tip. Time zero is the peak of actin assembly in the two-ring model. Time dependencies for pushing force (left), elongation speed (middle), and tubule length (right) are shown for rigid elongating invaginations with fixed threshold corresponding to turgor pressure 8 atm.

## Discussion

Endocytosis in fission and budding yeast depends on forces produced by the assembly of expanding networks of actin filaments, which drive invagination of the plasma membrane against the high internal turgor pressure. However, it was unclear whether actin assembly generates forces sufficient to overcome the turgor pressure.

We formulated a mathematical model of the forces based on principles of polymer physics that integrates the kinetics of the biochemical reactions (actin filament nucleation, elongation, capping, and severing), the rheological properties of actin filament networks and the time course of numbers of participating proteins. Certain modeling assumptions and approximations used in this study are similar to those adopted in other models of endocytosis in yeast (Carlsson and Bayly, 2014; Carlsson, 2018; Lacy et al., 2018; Mund et al., 2018). In particular, as in previous studies, we assume that movement is transmitted from a growing actin patch to the endocytic invagination via connections of actin filaments to the plasma membrane. As assumed previously (Carlsson and Bayly, 2014), our model approximates a network of actin filaments as a continuous medium, though Carlsson and coworkers (as well as the authors of a discrete model in (Mund et al. 2018)) approximate the actin patch as a growing elastic solid. Taking into account the turnover of actin in the patch, largely due to severing of the filaments by cofilin, we interpret the mechanics of endocytic actin meshwork as that of a viscoelastic fluid, with parameters constrained by measured rheological properties of overlapping filaments. This has yielded forces sufficient to withstand turgor pressure in fission yeast. Simulations of the model also reproduce the temporal and spatial distributions of actin at sites of endocytosis and explain the flask-shaped invaginations of the plasma membrane observed by electron microscopy (Kukulski et al., 2012).

Our model allows for different assumptions about the location of the nucleation-promoting factors that activate Arp2l3 complex to drive the assembly of the actin filament networks. We compared a two-ring hypothesis proposed for fission yeast (Arasada and Pollard, 2011; Arasada et al., 2018), a model proposed for budding yeast (Picco et al., 2015; Sun et al., 2017; Mund et al., 2018) where all NPFs remain at the base of the invagination, and a hypothetical model where the NPFs move with the tip of the invagination.

Simulations of the two-ring model produced two interacting zones of actin filaments with high densities near the rings. The internal repulsive stress generated by actin polymerization causes the entire patch to expand. Constraints imposed by the plasma membrane and cell wall result in expansion of the network inward and laterally, exerting drag on an initial invagination and thus pulling it inward. Given the known number of polymerized actin subunits and viscosity of the actin meshwork, we estimate the magnitude of this drag. The dendritic structure of the meshwork produces entanglement that enhances viscosity to levels sufficient to produce forces in the range of 2,200-3,000 pN, which, for invaginations with typical diameters, would overcome turgor pressure ~ 8-10 atm. The estimates are within the energy and critical load constraints, with the buckling threshold being met, in part, with the aid of crosslinking by fimbrin.

Simulations of the one-zone models with the numbers of nucleation-promoting factors and initial conditions used for the two-zone model also produced drag on the invagination. The budding yeast model with the NPFs remaining at the base of the invagination generated forces close to the two-ring model. This result underscores the importance of the cell wall to provide support for the actin filament network to generate a traction force. In the two-ring model mutual repulsion of the two zones of actin filaments compensates for the partial loss of support from the cell wall. The model with the NPFs moving at the tip generated significantly weaker forces, resulting in a much shorter invagination than the two other models.

The general model allowed us to simulate the forces required to elongate an endocytic tubule, although we used the simplifying assumption that the invagination is a (sphero)cylinder with a fixed radius. We also assumed that once the generated force overcomes the turgor threshold, all the points on the invagination move with the same (but time-dependent) speed *u*(*t*) = *μ*(*f*_push_(*t*) − *f_c_*). Somewhat counterintuitively, the speed and the length attained by the invagination is virtually insensitive to the mobility coefficient *μ*, but rather depends on how early during patch assembly the force produced by actin assembly *f*_push_(*t*) overcomes the opposing force from turgor pressure *f_c_*. For *f_c_* corresponding to the 8-atm turgor pressure, the simulations yielded a maximum tubule length somewhat shorter than experimental patch sizes.

We discovered that expansion of the actin filament network produces radial forces normal to the tubule. The distribution of these radial forces along the tubule will squeeze the invagination near its opening and stretch the middle, producing a shape like a flask as observed by electron microscopy in budding yeast (Kukulski et al., 2012). Without reliable information about elastic properties of endocytic invaginations we could not solve for shape of the invagination. However, a small pore between the exterior and the lumen of the invagination reduces *f_c_* as actin assembles. We approximated the effect of this shape change with a model with a decreasing threshold *f_c_* (*t*) over time to show that reducing the size of the pore favors the formation of longer tubules.

## Acknowledgements

Research reported in this publication was supported by National Institute of General Medical Sciences of the National Institutes of Health under award numbers R01GM026338, P41GM103313 and R01GM115636. The content is solely the responsibility of the authors and does not necessarily represent the official views of the National Institutes of Health. The authors thank Rajesh Arasada for advice and super resolution measurements of actin patch dynamics. M.N. and B.M.S. thank Leslie Loew for continuing support and helpful discussions.

## SUPPLEMENTAL MATERIAL

### Supplemental figures

**Supplemental Figure 1.**
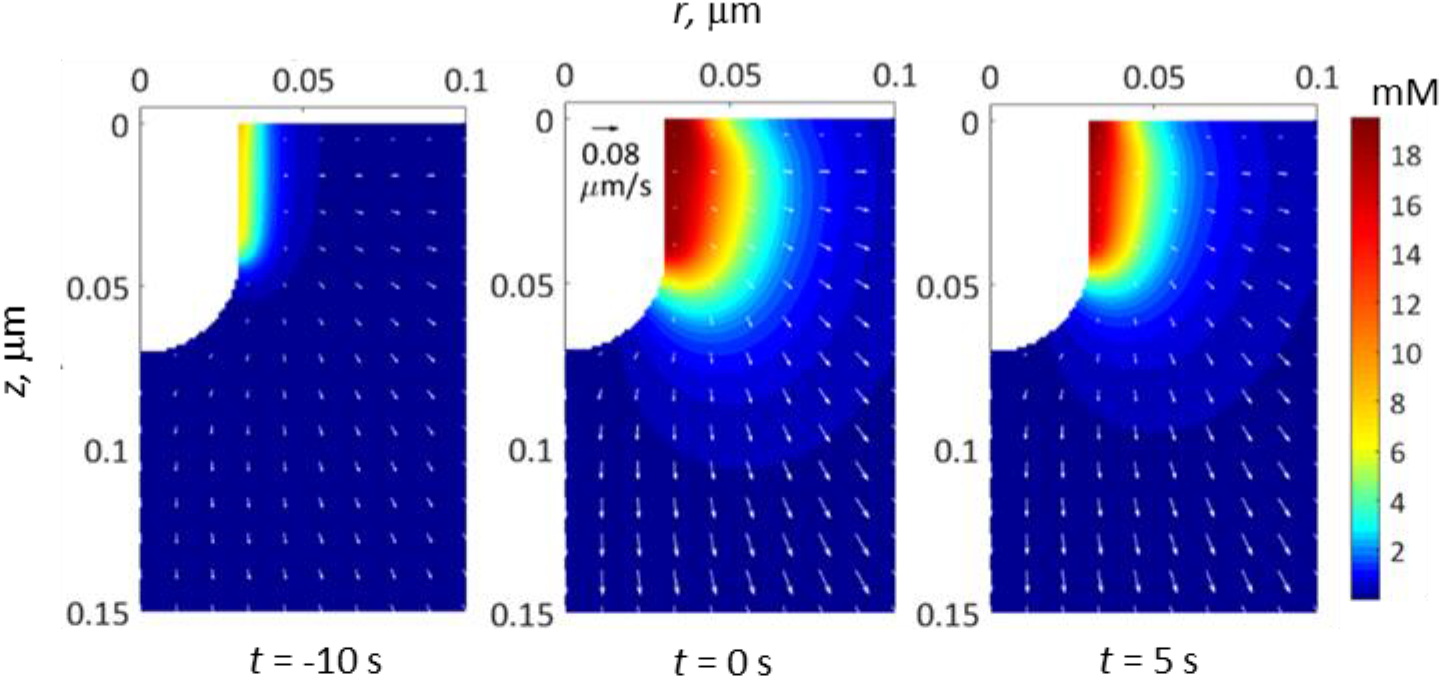
Simulation of a single ring model of actin patch assembly around a tubule with fixed geometry. Actin densities (pseudo-color) and velocities (arrows) are shown for *r-z* sections of 3D geometry and selected time points. Two rings of nucleation-promoting factors, not shown explicitly, were positioned next to each other at the base of the invagination adjacent to the horizontal portion of the plasma membrane.

**Supplemental Figure 2.**
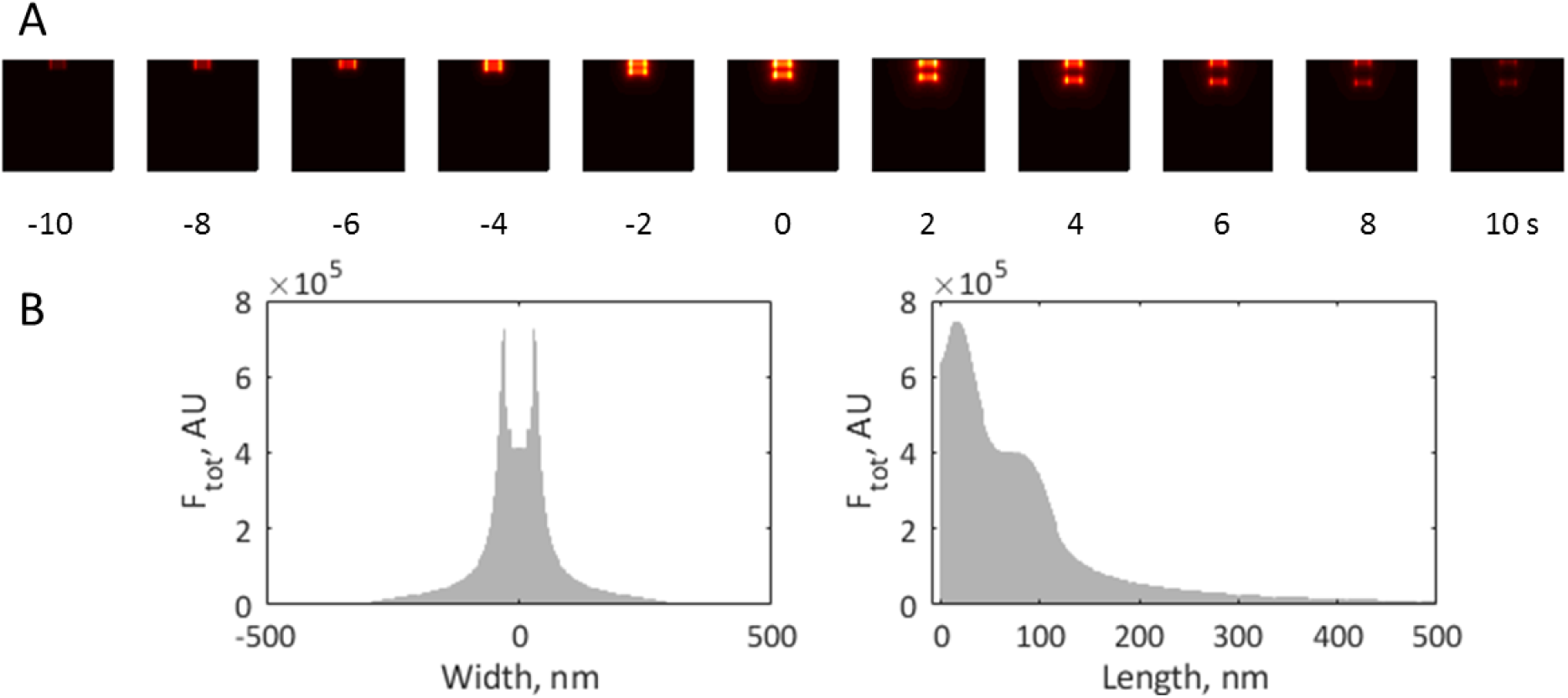
Results of simulating an elongating tubule with time-dependent force threshold, before applying median filter. (A) Heat maps of actin density (see Fig. 7C of main text) projected on plane of image are shown for selected time points. (B) Width and length distribution of actin density were obtained by integrating results of panel (A) over time. The trough of the width distribution reflects the existence of the space inside the invagination that is void of actin. No such troughs were observed experimentally, likely because the spatial resolution was comparable to the invagination width.

### Supplemental movies

**Supplemental Movie 1.** Simulation of elongating invagination with fixed resisting force.

Results are shown for turgor pressure 8 atm. Left panel: 3D distribution of actin density (pseudo-color) in the vicinity of the tubule. Middle panel: actin density and its velocities (arrows) are shown for *r-z* sections of 3D geometry. Right panel: deformable computational mesh used in simulation; deformations of mesh conform to moving invagination. Note that the movie was made with a color code built in COMSOL, which is slightly different from that used in the Figures.

**Supplemental Movie 2.** Simulation of elongating invagination with force threshold decreasing with time.

Left panel: 3D distribution of actin density (pseudo-color) in the vicinity of the tubule. Middle panel: actin density and its velocities (arrows) are shown for *r-z* sections of 3D geometry. Right panel: deformable computational mesh used in simulation; deformations of mesh conform to moving invagination. Note that the movie was made with a color code built in COMSOL, which is slightly different from that used in the Figures.

## Methods

### M.1 Governing Equations

#### M1.1 Computational Domain

Based on the assumptions described in *Model*, the computational domain depicted in Figure 1 of the main text remains axisymmetric throughout the elongation process. Because the localization of membrane-bound species and the corresponding fluxes are also axisymmetric, solutions of the model will have the same symmetry. Therefore, the problem reduces to solving an equivalent 2D model in cylindrical coordinates ***x*** = (*r, z*) in the domain *Ω* shown in Figure M1, where *Γ_i_*(*i* = 1, …, 5) are the corresponding boundaries. Note that the full 3D geometry is restored by revolving *Ω* around the axis of symmetry *r* = 0 (red dash-dotted line in Figure M1).

**Figure M1.**
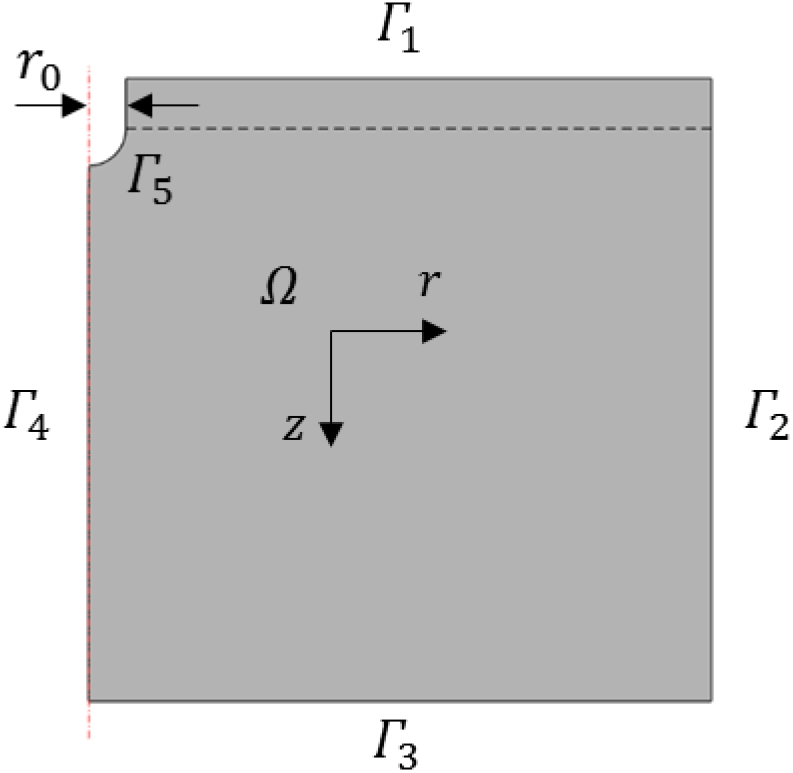
Equivalent 2D axisymmetric computational domain (*Ω*) and boundaries (*Γ*_1_, *Γ*_2_, *Γ*_3_, *Γ*_4_, *Γ*_5_).

The domain extensions (0.5 μm. in each coordinate direction) were chosen to be sufficiently large to ensure that numerical solutions are essentially independent from boundary conditions at *Γ*_2_ and *Γ*_3_ (see the following subsections). The cylindrical and hemispherical parts of the invagination degenerate in the 2D model into a line and a quarter of a circle, respectively. The initial length of the cylindrical part is 40 nm, as it accommodates two rings of nucleation promoting factors (NPFs), each being 20 nm wide (Arasada and Pollard, 2011). The radius of the endocytic invagination is *r*_0_ = 30 nm.

#### M1.2 Transport and Reaction Equations

Spatiotemporal dynamics of proteins involved in patch assembly are governed by conservation of mass, which in our model has the following form,

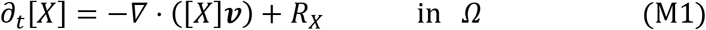

for all cytosolic species, except ActiveArp (the equation for ActiveArp is discussed below). In Eq (M1), [*X*] is the concentration of protein *X* in μM, *R_X_* is the sum of rates of all reactions affecting *X*, **v** is actin velocity, and *Ω* is the computational domain. In what follows, the density of actin network is defined as 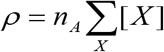, where the sum is taken over all cytosolic species, except for FArp and ActiveArp, and *n_A_* = 602 molecules μm^−3^/μM.

Functional forms of *R_X_* and parameters are taken from (Berro et al., 2010), with the exception of the on- and off-rate constant of polymerization, capping, and cofilin binding. The latter were modified by the factor (1 - *ρ*/*ρ*_max_)^*λ*^ that accounts for the effect of excluded volume. This factor ensures that the abovementioned processes slow down as *ρ* approaches *ρ*_max_, so that *ρ* never exceeds *ρ*_max_ = (4*πδ*^3^/3)^−1^, where *δ* = 2.7 nm is the subunit radius. The functional form used follows from the dependence of molecular diffusivities on the excluded volume (Novak et al., 2011). The parameters used in computations are *λ* = 0.5 (Novak et al., 2009) and *ρ*_mx_ = *n_A_* · 19.5 mM. The equations describing spatiotemporal dynamics of each species are listed below:

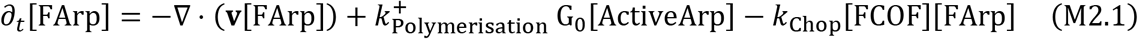

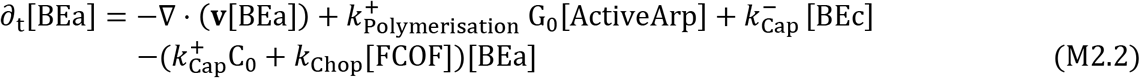

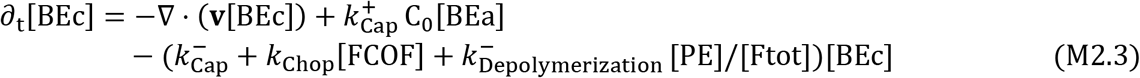

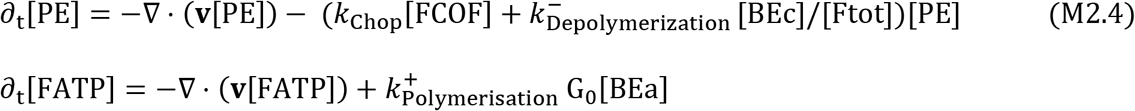

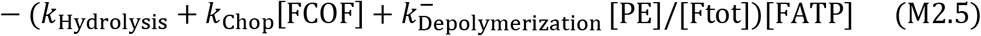

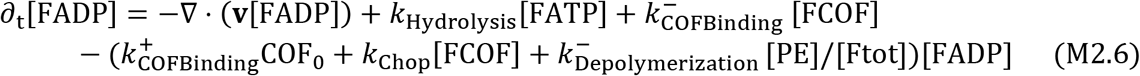

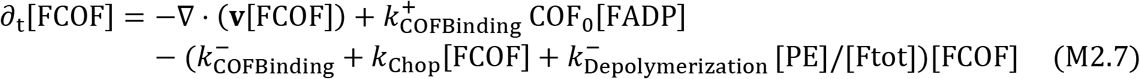

In Equations (M2.1) – (M2.7), Ftot = [FADP] + [FATP] + [FCOF] + [PE] + [BEa] + [BEc], and subscript ‘0’ denotes a constant. Values of reaction rate constants and constant concentrations are taken from Table 1 and Table 2 of (Berro et al., 2010). Equations (M2.1) – (M2.7) are subject to zero-flux boundary conditions at the membrane, 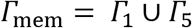, as well as at the boundary passing along the axis of symmetry *Γ*_4_ in Figure M1. Outflow boundary conditions were enforced at *Γ*_2_ and *Γ*_3_. Note that for solving equations (M2.1) – (M2.7), which are of the hyperbolic type, boundary conditions need not be specified on all *Γ_i_* (Ferziger and Perić, 2002). However, for technical reasons discussed in subsection *Finite Element Implementation of the Model*, a diffusion term with a very small diffusion coefficient was added to all equations. The resulting parabolic equations require boundary conditions on all boundaries of the domain.

We now describe the equation for ActiveArp. Active Arp2l3 complexes appear in the cytosol due to the flux of FArpTernCompl that only exists at the NPF rings (Figure 1 of the main text). The corresponding flux density is 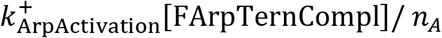, where [FArpTernCompl] is in moleculeslμm^2^. Because the detachment of FArpTernCompl from the membrane involves diffusion, a consistent description of [ActiveArp] near the rings should include a diffusion term. Therefore, the dynamics of [ActiveArp] is described by a diffusion-advection-reaction equation,

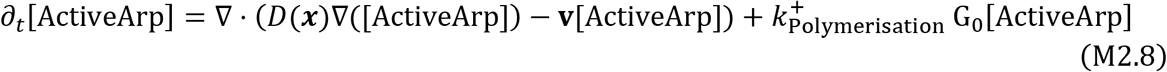

and a corresponding boundary condition,

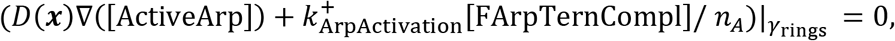

where *γ*_rings_ are the fragments of *Γ*_mem_ occupied by the rings. The diffusion term is restricted to the vicinity of the rings, by using a diffusion coefficient that is non-zero only along the cylindrical part of the tubule (*Γ*_5_ in Figure M1) and decays exponentially in the radial direction,

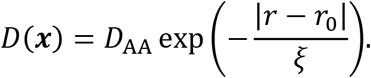

The parameter values used in the solutions were *ξ* = 3 nm and *D*_AA_ = 0.001 μm^2^/s, though changing the diffusion constant *D*_AA_ by several orders of magnitude did not change the outcome in any significant way. At all other boundaries, the conditions for [ActiveArp] were the same as for the other cytosolic species.

As in (Berro et al., 2010), adapter proteins that recruit and activate NPFs are not included in our model. Instead, a temporal wave of NPFs with a Gaussian shape drives actin assembly near the rings. Therefore, the surface densities of the membrane-bound proteins are governed by ordinary differential equations (ODEs) based on the rates of corresponding biochemical reactions:

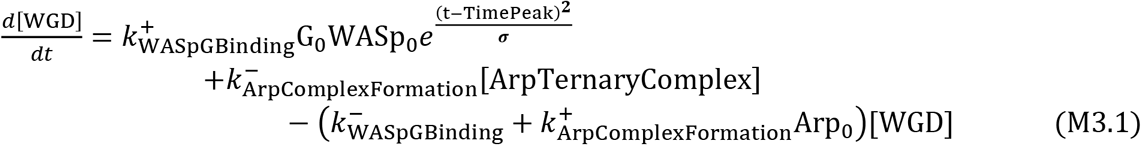

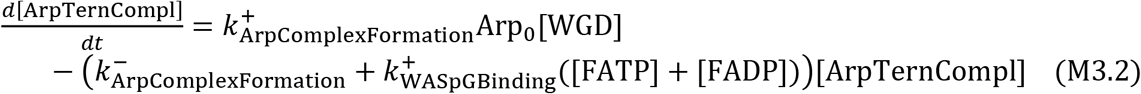

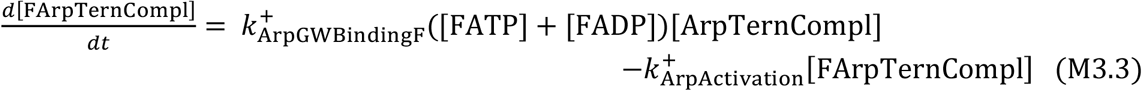

The densities of the membrane-bound species are in 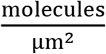. The reaction rates, initial conditions and other constants are taken from Table 1 and Table 2 of (Berro et al., 2010); note that the value of WASP_0_ was converted from μM to 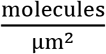.

#### M1.3 Actin Meshwork Mechanics Equations

The actin meshwork is modeled as a compressible visco-active fluid. In a viscosity-dominated environment of the actin patch, forces due to the fluid’s inertia and acceleration are neglected, which leads to a quasistatic formulation of the meshwork velocities **v**

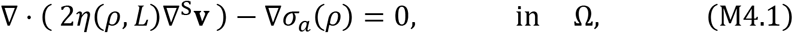

where 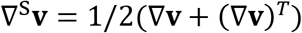 is the symmetrized velocity gradient tensor, *η*(*ρ,L*) = *κ*_visc_*ρ*(1/*N* + *ρδ*^2^*L*) is the dynamic viscosity, and *σ_a_* = *κ*_active_*ρ*^2^ is the active stress. See subsection *Model* for further details regarding the derivation of the functional forms of the viscosity and the active stress.

Equations (M4.1) are elliptic in nature, similar to the Stokes equations of a Newtonian fluid, and hence require boundary conditions on all boundaries of the computational domain. No-slip boundary condition is applied where actin meshwork meets the membrane

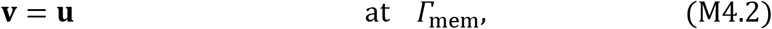

where **u** is the velocity of the membrane. All other boundaries are subject to zero-stress boundary conditions,

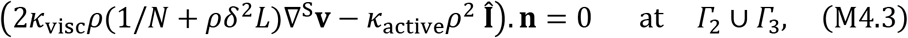

where 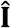 is a unit tensor and **n** is the outward normal vector to the boundary.

#### M1.4 M1.4 Boundary Conditions and Domain Size Effects

The simulations were first run in a domain with smaller extensions in each coordinate direction, 0.3 μm instead of 0.5 μm. To ensure that the boundary conditions applied at *Γ*_2_ and *Γ*_3_ had no effect on the numerical results, we ran simulations with different types of boundary conditions and in larger domains. No significant changes in the solutions were observed. All the numerical results presented in the paper are from the simulations performed in the larger domains, 0.5 μm in each coordinate direction.

### M2 Moving Boundaries Formulation

#### M2.1 Modelling Tubule Movement

Simulations of elongating invaginations involve additional assumptions. In particular, the shape of the invagination is assumed to remain (sphero)cylindrical during the elongation process, so that only the cylindrical part elongates. Furthermore, we assume for simplicity that the invagination is infinitely rigid, so that all material points move with a same instantaneous velocity, which changes linearly with the net viscous drag exerted by the actin network; the linear dependence on the pushing force is parameterized by a mobility coefficient, see Equation (5) in *Model*.

From fluid mechanics, the viscous forces acting on the tubule are given by the integral of the total stress in the actin meshwork over the surface of the endocytic invagination,

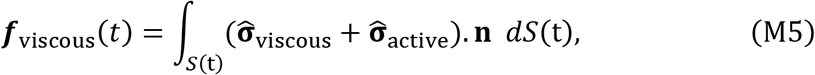

In Eq (M5), integration is carried over the time-dependent boundary S(*t*) = *Γ*_5_(*t*) representing the invagination, and **n** = (**n**_*r*_, **n**_*z*_)^*T*^ is the outward unit normal vector to the boundary *Γ*_5_(*t*) (directed from *Γ*_5_(*t*) towards the interior of *Ω*(*t*)). The velocity of the tubule at any given time is then obtained by Eq (5) of *Model*. The *z*-component of the viscous force is the drag force exerted on the invagination, *f*_z_(*t*) = **f**_viscous_(t) · ***n**_z_*, and the force due to the turgor pressure *Π*_turgor_, is *f*_c_ = 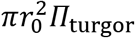, where *r*_0_ is the radius of the (sphero)cylindrical invagination (Figure M1); if the invagination is constricted by the surrounding meshwork, *r*_0_ is the radius of the pore between the exterior and the lumen of the invagination.

#### M2.2 The ALE Framework

The models of elongating invaginations were solved using an Arbitrary Lagrangian-Eulerian (ALE) method. The ALE method is described in numerous publications, see e.g. (Donea et al., 2004). In an ALE simulation, the computational mesh moves with displacementslvelocities prescribed at the boundaries of interest (normally loading and interface boundaries). At all other places in the domain, the mesh moves with a smooth arbitrary velocity such that mesh quality is maintained throughout the simulation, while mesh connectivity remains the same. The governing equations formulated in a Eulerian coordinate system should be reformulated based on the ALE framework. Following the notation used by (Formaggia and Nobile, 2004), a fixed reference frame i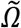 and a mapping : 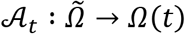 is defined to provide a one-to-one correspondence 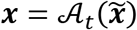, and 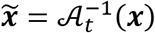 between the Eulerian coordinates ***x*** = (*r, z*) ∈ Ω(*t*) and ALE coordinates 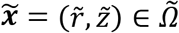. It is straightforward to show that for any scalar function *f*(***x**, t*), the Eulerian and ALE time derivatives are related by the chain rule,

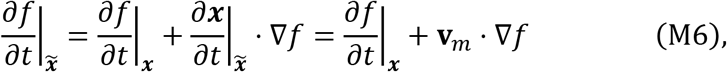

where 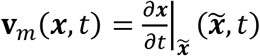 is the local mesh velocity. The mesh velocity can be obtained by solving in the domain a mesh smoothing equation. See *Finite Element Implementation of the Model* for further details.

Since domain *Ω*(*t*) changes with time, it is generally not possible to discretize directly the Eulerian time derivatives in the transport-reaction equations. In fact, if ***x*** ∈ *Ω*(*t*) and Δ*t* > 0, the condition ***x*** ∈ *Ω*(*t* + Δ*t*) may not be always satisfied (San Martín et al. 2009). Therefore, the Eulerian time derivatives 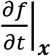 in the transport-reaction equations are substituted by the right-hand side of equation (M6). This introduces additional advection-like terms to the equations with the advection velocity being the local mesh velocity **v**_*m*_. For example, the transport equation (M1) in the equivalent ALE formulation reads as

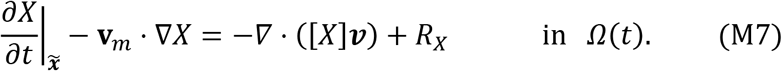

It should be noted that all space derivatives in Equation (M7) are taken with respect to the Eulerian coordinates ***x*** This equation is subject to Rankine-Hugoniot boundary condition (zero-flux boundary condition) on the moving boundary *Γ*_5_(*t*). Boundary conditions on all other non-moving boundaries remain unchanged.

The equations for actin meshwork mechanics and their boundary conditions do not change in the ALE framework. This is because these equations are in quasistatic form and there are no history-dependent rates in the definitions of viscous and active stresses (Donea et al., 2004).

ODEs that govern membrane-bound species are not modified as a result of the movement, since these species are treated in the model as non-spatial.

#### M2.3 Movement of the NPF Ring(s)

According to the two-ring hypothesis (Arasada and Pollard, 2011), two NPF rings drive the actin assembly. One of the rings remains stationary near the horizontal membrane, *Γ*_1_ in Figure M1. The other ring moves with the tubule, keeping its proximity to the tip of the tubule. During the movement the width of the NPF rings and their radius remain constant. Therefore, it suffices to track the z-component of the position of the moving ring *z*_ring_ described by

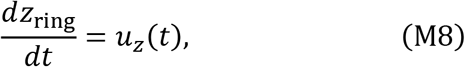

where *u_z_*(*t*) is as in Eq (5) of *Model*. The movements of the rings were tracked similarly in the one-ring models described in *Results*.

### M3 Finite Element Implementation of the Model

We used a Galerkin finite-element method to solve numerically the governing equations for the transport and reaction of proteins, and the equation for velocities of the actin meshwork. These equations are implemented and solved in COMSOL Multiphysics (COMSOL, 2015) in a 2D axisymmetric domain (Figure M1), as described below.

#### M3.1 Computational Mesh

A computational mesh used for spatial discretization of the governing equations consisted of 33777 quadrilateral elements (Figure M2(a)). To approximate the velocity gradients near the invagination with more precision, a boundary layer mesh was constructed. These gradients are important for calculating forces exerted on the tubule, and they affect the accuracy of the numerical solution overall. Figure M2(b) is a zoomed-in view of the vicinity of the invagination to show the boundary layer mesh.

**Figure M2:**
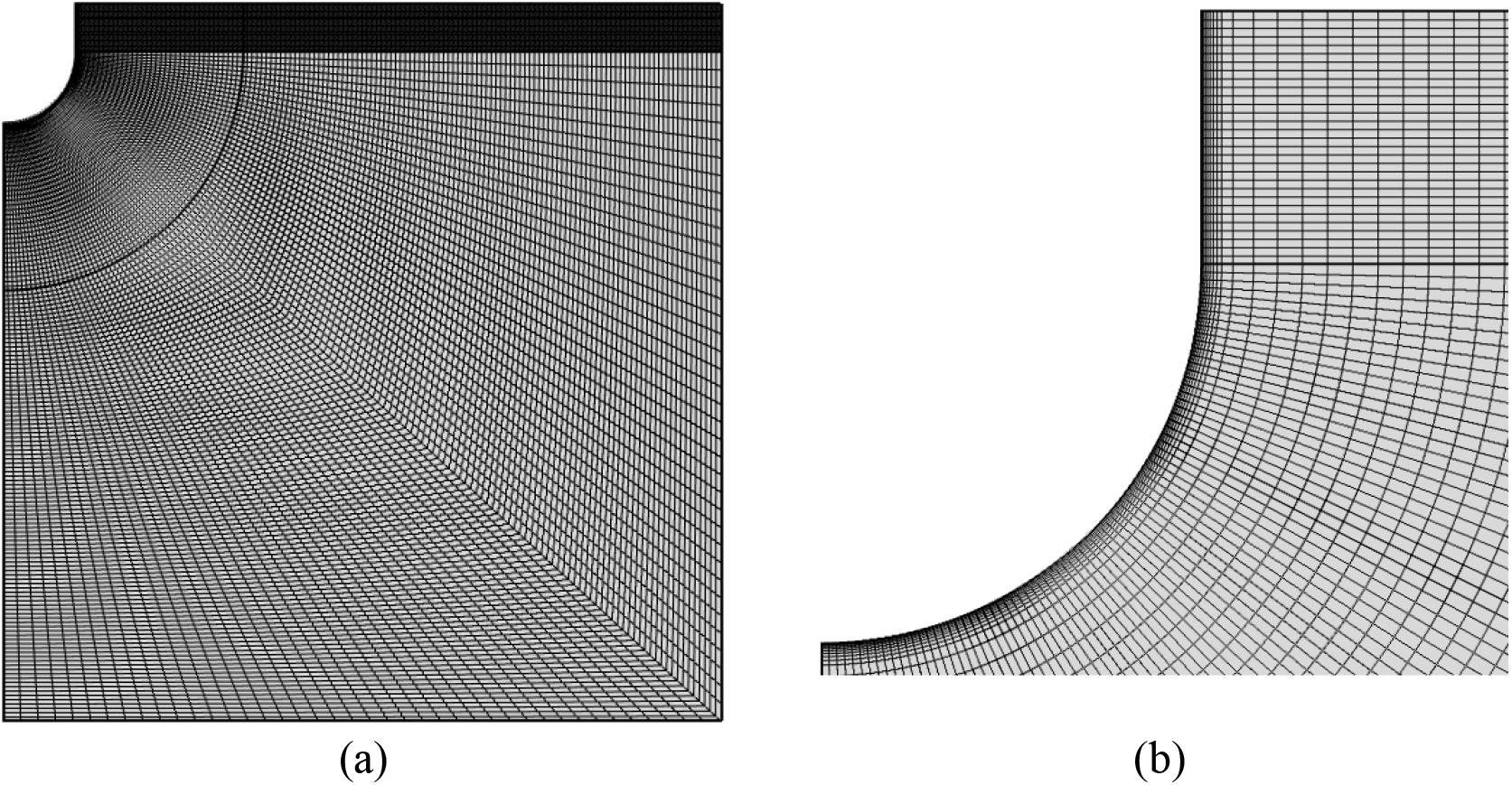
The computational mesh(a), and a zoomed-in view near the invagination boundaries (b).

The mesh was designed so that as the tubule grew, the elements near the horizontal membrane and in the vicinity of the cylindrical part of the tubule were elongated in the z direction. To maintain sufficiently fine elements even after they were stretched as a result of the elongation, a high initial mesh density was used in the vertical direction in these regions. For more details about the design of mesh movements and its implementation see subsection *Mesh Smoothing Equations* below.

Classical mesh refinement was performed for simulations in fixed geometries and for one simulation of an elongating invagination to ensure that numerical results were grid-independent. The original mesh was refined by reducing the linear size of elements by approximately a factor of 2. This yielded 132884 quadrilateral elements, roughly four times the number of elements in the original mesh. The solutions obtained with refined meshes differed from the original mesh by less than 0.3%.

Given the negligible differences, all subsequent moving geometry simulations were performed on the original mesh.

#### M3.2 Transport Equations of Cytosolic Species

Eqs (M2.1-M2.8), governing the spatiotemporal dynamics of cytosolic species, were solved using COMSOL’s ‘Transport of Diluted Species’ module. For simulating moving domains, the module automatically adds to the transport equations advection-like terms of Eq (M7). Linear Lagrange finite elements were used to approximate the concentrations of these species.

Solving Eqs (M2.1-M2.7) with the standard Galerkin finite-element method may result in spurious oscillations (Donea and Huerta, 2003). Treating advection terms with Petrov-Galerkin type methods available in COMSOL can suppress these unphysical oscillations. However, the effectiveness of these methods generally depends on values of auxiliary parameters, and some numerical oscillations may persist. We chose instead adding to the transport equations a diffusion term with a small diffusion coefficient, termed ‘technical diffusion’ with diffusivity *D*_tech_., and using standard discretization schemes for all terms. For consistency, *D*_tech_. was added also to *D*(***x***) in Eq (M2.8) for [ActiveArp]. In all simulations, we used the value *D*_tech_. = 1 × 10^−5^μm^2^/*s*. Decreasing *D*_tech_. further by an order of magnitude did not produce significant changes in the solution. While spurious oscillations can occur in solving diffusion-advection equations on meshes with high Peclet numbers, no such oscillations were observed after adding technical diffusion for the meshes used in our computations (see subsection M3.1).

#### M3.3 ODEs for Membrane-bound Species

Eqs (M3.1-M3.3) for membrane-bound species were solved on *Γ*_5_ (Figure M1) with the ‘Boundary ODEs and DAEs’ module of COMSOL. Positions of the rings of NPFs were accounted for by multiplying the first term in the right-hand side of Eq (M3.1) by a Boolean expression, which was evaluated to one at the locations of the rings and zero elsewhere. As the rings moved, the expression was updated accordingly. Although, membrane-bound species are non-zero only at the locations of the rings, the corresponding ODEs were solved everywhere on *Γ*_5_, allowing for a uniform application of the flux boundary condition for [ActiveArp], although the flux density was non-zero only at γ_rings_. Constant discontinuous Lagrange finite elements were used for the membrane-bound species.

#### M3.4 Velocity Equations

Eqs (M4.1) for actin velocities were solved using the ‘Weak Form PDE’ module of COMSOL, which allows one to implement a method of weighted residuals solving equations in weak forms (Donea and Huerta, 2003). Let 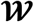 be the space of weighting (test) functions vanishing on the Dirichlet boundaries *Γ*_mem_, and let 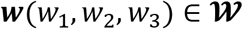 be the test functions for velocities in the cylindrical coordinates. The weighted residual form of equations in the moving domain *Ω*(*t*) is then written as

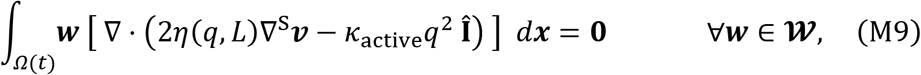

where *q* and *υ* are the weak solutions corresponding to the polymerized actin density *ρ* and actin velocities **v**. The weak solution *υ* resides in a space of admissible functions satisfying the Dirichlet (no-slip) boundary condition (M4.2). Integrating by parts and applying Green’s formula (Donea and Huerta, 2003) then yields

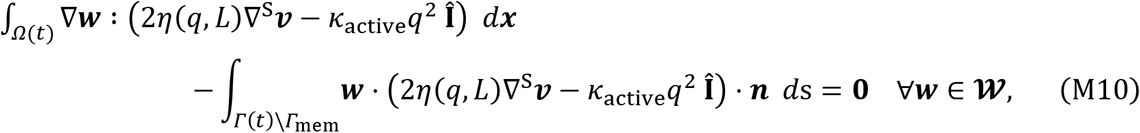

where ‘:’ denotes the double dot product of two tensors. The integrand in the second integral of equation (M10) is zero on *Γ*(*t*)\*Γ*_mem_ due to the zero-stress boundary condition (M4.3), so the final weak form of the velocity equations reads

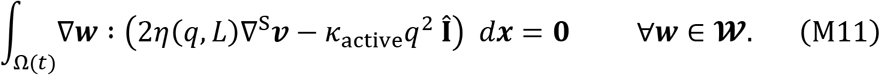

To derive equations for velocity components in weak form for the equivalent two-dimensional axisymmetric coordinate system, one should start with the full differential operators in cylindrical coordinates (*r, θ, z*), and then remove *θ*-components and derivatives with respect to *θ*. In a cylindrical coordinate system with orthonormal basis vectors 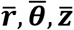, the velocity gradient operator and the symmetrized velocity gradient tensor applied to the weak solution *υ* = (*υ*_1_, *υ*_2_, *υ*_3_)^*T*^ are defined as follows:

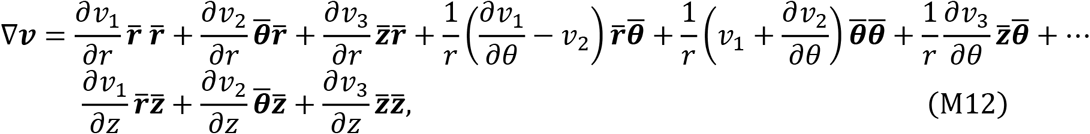

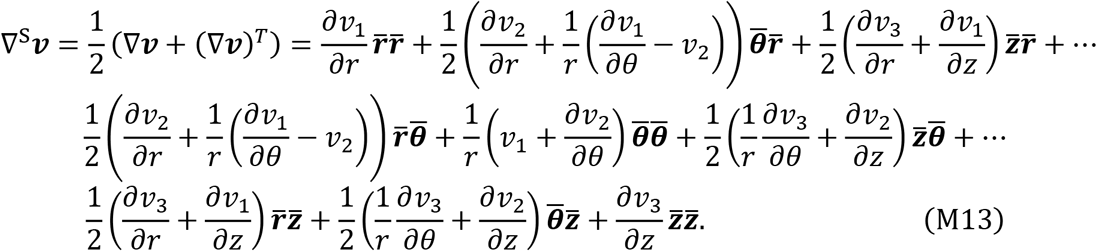

A unit tensor is defined as 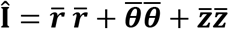. Using these definitions, the first term in the integrand of equation (M11), 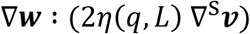, is

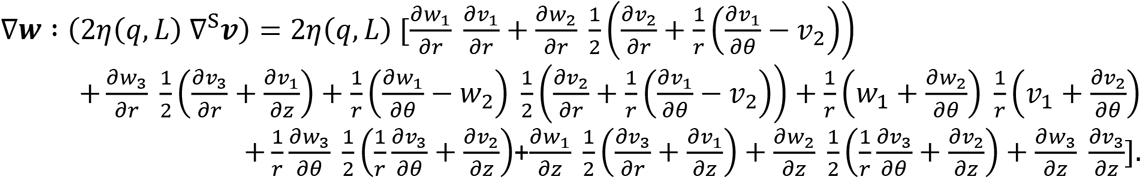

The simplification due to axial symmetry yields the following weak form of the first term in Eq (M11):

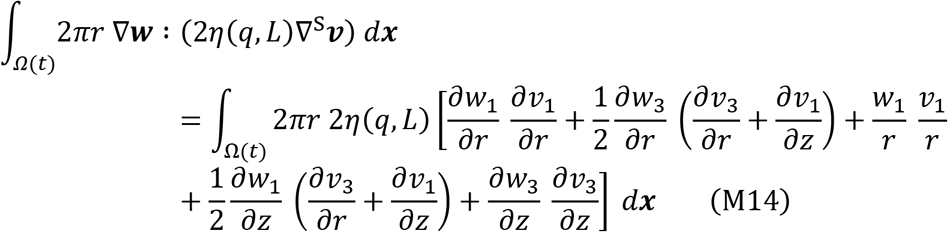

Similarly, the second term of the integrand in Eq (M11) yields

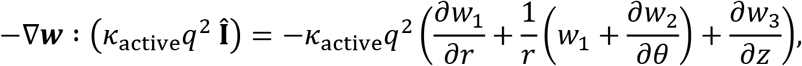

and upon the reduction due to axial symmetry, the weak form of the second term in Eq (M11) is

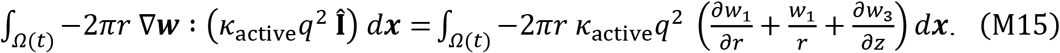

The factor 2*πr* in Eqs (M14-15) is the result of integration over *θ*

Eqs (M14) and (M15) were implemented in COMSOL. Linear Lagrange finite elements were used in computing actin velocities.

#### M3.5 Mesh Smoothing Equations

Solving a moving boundary problem using the ALE method requires computing local mesh velocities **v**_m_. While **v**_m_ are not known in advance in the interior of the domain, velocities of points on a moving tubule are computed from Eq (5) of *Model*, while other boundaries of the computational domain are fixed in the course of a simulation. To correctly model the movements of the domain, mesh velocities at the boundaries should coincide with the velocities of the boundary. Then the mesh velocities of the interior points of the domain may be computed, for instance, by employing a harmonic extension of the boundary velocities (Formaggia and Nobile, 2004).

Computing **v**_m_ and tracking of mesh movements were done using the ‘Moving Mesh’ module of COMSOL, which allows one to prescribe mesh displacements **x**_m_ and/or mesh velocities at the domain boundaries and at any other interior domain points/edges. Values of **v**_*m*_ in the domain interior were computed using a Laplacian mesh smoother with linear geometric shape functions. Care must be exercised in simulating large elongations, which may result in a highly distorted mesh. The ALE methods become instable on distorted meshes, so that the domain needs to be remeshed to restore the regularity of the elements (San Martín et al., 2009). Remeshing entails interpolation to a new mesh, which introduces additional error. Also, frequent remeshing increases computational costs. To avoid remeshing and the issues associated with it, we defined a virtual edge in the interior of the computational domain, indicated by a dashed line in Figure M1. The tubule velocity computed from Eq (5) in *Model* was then used as the z-component of the mesh velocity for both the virtual edge and the circular part of *Γ*_5_. The r-components of the mesh velocity on these segments were set to zero. The prescribed movement of the virtual edge guides the mesh deformation in the interior of the domain and allows for modeling very large tubule elongations without remeshing. On *Γ*_2_, *Γ*_4_ and the straight part of *Γ*_5_, the r-component of the mesh displacement was set to zero, whereas the vertical components was allowed to vary freely. Displacements of the mesh on the remaining horizontal segments of the domain boundary were set to zero.

#### M3.6 Equation for Updating NPF Ring Position

Eq (M8), determining the time-dependent z-component of the position of the moving ring, *z*_ring_, was solved using COMSOL’s ‘Point ODEs and DAEs’ module for one point on *Γ*_5_. The *z*_ring_ was initialized to the position of the ring at *t* = 0. Because Eq (M8) was solved in COMSOL within a spatial model, a constant discontinuous Lagrange finite element was used to approximate *z*_ring_.

### M4 Solvers and Computational Parameters

The coupled nonlinear system of equations describing the cytosolic species, Eqs (M2.1-8), the membrane-bound species, Eqs (M3.1-3), the ring’s position Eq (M8), and the actin velocities, Eqs (M14-15), along with the corresponding boundary conditions, were discretized using FEM and solved in a fully coupled manner in COMSOL. Note that even though the force-balance equation does not involve time derivatives, the coupled system constitutes an initial-value problem, so that initial conditions must be specified for all variables (initial values of the actin velocities were set to zero).

The time-dependent system was solved using a backward-differentiation time-stepping method of order 1-2. Relative and absolute tolerances of the time-stepper were set to 1 × 10^−5^ and 1 × 10^−6^, respectively. Other default solver parameters were used without modification. Linearization was performed using Newton’s method with a constant damping factor of 1. The system’s Jacobian was updated at each nonlinear iteration. The linearized system was solved monolithically using a direct MUMPS solver with default solver parameters. We verified, by solving the problem with varying solver parameters (including the tolerances of the time-stepper), that the solutions did not depend on specific choices of parameters of the solver.

### M5 Data Analysis and Display

Presentation and post-processing of numerical results were facilitated by exporting the COMSOL FEM solutions, obtained at the Lagrange points, which were further processed in MATLAB R2017b [ref(s)]. The 2D snapshots of the solution (see, as an example, Figure 4 of the main text or in Supplemental Figures 1) were obtained by interpolating the FEM solutions onto a uniform 2D grid. A sufficiently large size of the grid allowed for accurately capturing all important features of the FEM solution that were first visualized in COMSOL. The 3D snapshots (see, for instance, Figure 7A of the main text or in Supplemental Movies 1 and 2) were exported as image files from COMSOL and then replotted in MATLAB.

The actin filament heat maps in Figure 8 and in the Supplemental Figure 2 were produced by first interpolating the FEM solutions for polymerized actin onto a uniform 3D grid defined inside a domain with the horizontal and vertical extensions of [–0.5, 0.5] μm and [0,0.5] μm, equal to the respective ranges of *r* and z coordinates of the 2D axisymmetric model. The extension in the depth direction was [–0.2,0.2] μm, in accordance with the thickness of the imaging plane in epifluorescence microscopy experiments of (Arasada et al., 2018). The interpolated 3D data was then projected on a 2D plane by integrating over the depth direction; the corresponding heat maps are presented in Supplemental Figure 2. The projected data were further subjected to a median filter with a half window size of 35 nm; the heat maps for the filtered data are shown in Figure 8 of the main text.

The histograms in Figure 8 (and in Supplemental Figure 2) were produced as follows. First, the filtered projected (projected only for Supplemental Figure 2) data were integrated over time. This yielded a two-dimensional matrix with the elements corresponding to the 2D image of the actin filament density integrated over time. In accordance with the protocol adopted by (Arasada et al., 2018), the width (length) distribution of the actin density was generated by summing up the values of the elements in each column (row) of the matrix. The width of the patch was calculated as the width of the corresponding histogram at half its maximum.

## References

Aghamohammadzadeh, S., and K. R. Ayscough. 2009. Differential requirements for actin during yeast and mammalian endocytosis. Nat. Cell Biol. 11:1039–1042.

Arasada, R., and T.D. Pollard. 2011. Distinct roles for F-BAR proteins Cdc15p and Bzz1p in actin polymerization at sites of endocytosis in fission yeast. Curr. Biol. 21:1450–1459

Arasada, R., W.A. Sayyad, J. Berro and T.D. Pollard. 2018. High-speed superresolution imaging of the proteins in fission yeast clathrin-mediated endocytic actin patches. Mol. Biol. Cell. 29:295–303.

Basu, R., E. L. Munteanu, and F. Chang. 2014. Role of turgor pressure in endocytosis in fission yeast. Mol. Biol. Cell. 25:679–687.

Berro, J., and T.D. Pollard. 2014. Synergies between Aip1p and capping protein subunits (Acp1p and Acp2p) in clathrin-mediated endocytosis and cell polarization in fission yeast. Mol. Biol. Cell 25:3515–3527.

Berro, J., V. Sirotkin, and T.D. Pollard. 2010. Mathematical modeling of endocytic actin patch kinetics in fission yeast: disassembly requires release of actin filament fragments. Mol. Biol. Cell. 21:2905–2915.

C.P. Broedersz., and F. C. MacKintosh. 2014. Modeling semiflexible polymer networks. Rev. Mod. Phys. 86:9951036.

Buxbaum, R.E., T. Dennerll, S. Weiss, and S.R. Heidemann. 1987. F-actin and microtubule suspensions as indeterminate fluids. Science 235:1511–1514.

Carlsson, A.E. 2018. Membrane bending by actin polymerization. Curr. Opin. Cell Biol. 50:1–7.

Carlsson, A. E., and P. V. Bayly. 2014. Force generation by endocytic Actin patches in budding yeast. Biophys. J. 106:1596–1606.

Chen, Q., and T. D. Pollard. 2013. Actin filament severing by cofilin dismantles actin patches and produces mother filaments for new patches. Curr. Biol. 23:1154–1162.

Cox, W.P., and E.H. Merz. 1958. Correlation of dynamic and steady-flow viscosities. J. Polym. Sci. 28:619–622.

Doi, M., and S. Edwards. 1998. The Theory of Polymer Dynamics, Oxford University Press.

Footer, M.s J., J. W.J. Kerssemakers, J. A. Theriot, and M. Dogterom. 2007. Direct measurement of force generation by actin filament polymerization using an optical trap. PNAS. 104:2181–2186.

Gardel, M.L., M.T. Valentine, J.C. Crocker, A.R. Bausch, and D.A. Weitz. 2003. Microrheology of entangled F-actin solutions. Phys. Rev. Lett. 91:158302.

Kaksonen M, and A. Roux. 2018. Mechanisms of clathrin-mediated endocytosis. Nat. Rev. Mol. Cell. Biol. 19:313–326.

Kasza, K.E., C.P. Broedersz, G.H. Koenderink, Y.C. Lin, W. Messner, E.A. Millman, F. Nakamura, T.P. Stossel, F.C. MacKintosh, and D.A. Weitz. 2010. Actin Filament Length Tunes Elasticity of Flexibly Cross-Linked Actin Networks. Biophys J. 99:1091–1100.

Kruse, K., J-F. Joanny, F. Jülicher, J. Prost, and K. Sekimoto. 2005. Generic theory of active polar gels: a paradigm for cytoskeletal dynamics. Eur. Phys. J. E 16:5–16.

Kukulski, W., M. Schorb, M. Kaksonen, and J. A. Briggs. 2012. Plasma membrane reshaping during endocytosis is revealed by time-resolved electron tomography. Cell 150:508–20.

Lacy, M.M., R. Ma, N.G. Ravindra, and J. Berro. 2018. Molecular mechanisms of force production in clathrin-mediated endocytosis FEBSLett. 592:3586–3605.

L.D. Landau, and E.M. Lifshitz. 1987. Fluid Mechanics: Volume 6 (Course of Theoretical Physics).

F.C. MacKintosh, J. Käs, and P. A. Janmey. 1995. Elasticity of Semiflexible Biopolymer Networks. Phys. Rev.Lett. 75:4425.

W.M. McFadden, P. M. McCall, M.L Gardel., and E. M. Munro. 2017. Filament turnover tunes both force generation and dissipation to control long-range flows in a model actomyosin cortex. PLOS Comp. Biol. 13(12): e1005811.

R.D Mullins., J. F. Kelleher, J. Xu, and T. D. Pollard. 1998. Arp2l3 complex from Acanthamoeba binds Profilin and cross-links Actin filaments. Mol. Biol. Cell 9:841–852.

Mund, M., J.A. van der Beek, J. Deschamps, S. Dmitrieff, P. Hoess, J-L. Monster, A. Picco, F. Nédélec, M. Kaksonen, and J. Ries. 2018. Systematic nanoscale analysis of endocytosis links efficient vesicle formation to patterned actin nucleation. Cell 174:884–896.

Nickaeen, M., I. L Novak, S. Pulford, A. Rumack, J. Brandon, B. M. Slepchenko, and A. Mogilner. 2017. A free-boundary model of a motile cell explains turning behavior. PLOS Comp. Biol. 13(11): e1005862.

Novak, I.L., F. Gao, P. Kraikivski, and B.M. Slepchenko. 2011. Diffusion amid random overlapping obstacles: Similarities, invariants, approximations. J. Chem. Phys. 134:154104.

Novak, I.L., P. Kraikivski, and B.M. Slepchenko. 2009. Diffusion in cytoplasm: effects of excluded volume due to internal membranes and cytoskeletal structures. Biophys J. 97:758–67.

Peskin, C. S., G. M. Odell, and G. F. Oster. 1993. Cellular motions and thermal fluctuations: the Brownian ratchet. Biophys. J. 65:316–324.

Picco, A., M. Mund, J. Ries, F. Nédélec, M. Kaksonen. 2015. Visualizing the functional architecture of the endocytic machinery. Elife. 4: e04535.

Prost, J., F. Jülicher, and J-F. Joanny. 2015. Active gel physics. Nat. Phys. 11: 111–117.

Satcher, R.L., Jr., and C.F. Dewey Jr., 1996. Theoretical estimates of mechanical properties of the endothelial cell cytoskeleton. Biophys. J. 71:109–118.

Sato, M., W. H. Schwarz, and T. D. Pollard. 1987. Dependence of the mechanical properties of actinlα-actinin gels on deformation rate. Nature 325:828–830.

Scher-Zagier, J. K., and A. E. Carlsson. 2016. Local turgor pressure reduction via channel clustering. Biophys. J. 111:2747–2756.

Sirotkin, V., J. Berro, K. Macmillan, L. Zhao, and T.D. Pollard. 2010. Quantitative analysis of the mechanism of endocytic actin patch assembly and disassembly in fission yeast. Mol. Biol. Cell 21:2894–2904.

Sun, Y., N. T. Leong, T. Jiang, A. Tangara, X. Darzacq, D.G. Drubin. 2017. Switch-like Arp2l3 activation upon WASP and WIP recruitment to an apparent threshold level by multivalent linker proteins in vivo. Elife 6: e29140.

Tseng, Y., and D. Wirtz. 2004. Dendritic branching and homogenization of Actin networks mediated by Arp2l3 complex. Phys. Rev. Lett. 93:258104.

Wachsstock, D.H., W.H. Schwarz, and T.D. Pollard. 1994. Cross-linker dynamics determine the mechanical properties of actin gels. Biophys. J., 66:801–809.

Wirtz, D. 2009. Particle-tracking microrheology of living cells: principles and applications. Annu. Rev. Biophys. 38:301–26.

Zaner, K.S., and T.P. Stossel. 1983. Physical basis of the rheologic properties of F-actin. J. Biol. Chem.258:11004–9.

## References

Arasada, R., W.A. Sayyad, J. Berro and T.D. Pollard. 2018. High-speed superresolution imaging of the proteins in fission yeast clathrin-mediated endocytic actin patches. Mol. Biol. Cell. 29: 295-303.

Arasada, R., and T.D. Pollard. 2011. Distinct Roles for F-BAR Proteins Cdc15p and Bzz1p in Actin Polymerization at Sites of Endocytosis in Fission Yeast. Curr. Biol. 21: 1450-1459.

Berro, J., V. Sirotkin, and T.D. Pollard. 2010. Mathematical Modeling of Endocytic Actin Patch Kinetics in Fission Yeast: Disassembly Requires Release of Actin Filament Fragments. Mol. Biol. Cell. 21: 2905-2915.

COMSOL Multiphysics. 2015. Version 5.2 [software]. Stockholm, Sweden: COMSOL AB. Available from: www.comsol.com.

Donea, J., and A. Huerta. 2003. Finite Element Methods for Flow Problems. Wiley.

Formaggia L., and F. Nobile. 2004. Stability analysis of second-order time accurate schemes for ALE-FEM. Comput. Methods Appl. Mech. Engrg. 193: 4097-4116.

Ferziger, J. H., and M. Perić. 2002. Computational Methods for Fluid Dynamics. Springer.

Donea, J., A. Huerta, J.-Ph. Ponthot, and A. Rodriguez-Ferran. Arbitrary Lagrangian-Eulerian Methods. 2004. In Stein, E., R. de Borst, and T.J.R. Hughes (eds) Encyclopedia of Computational Mechanics. Volume 1: Fundamentals. John Wiley & Sons.

Novak, I.L., P. Kraikivski, and B.M. Slepchenko. 2009. Diffusion in cytoplasm: effects of excluded volume due to internal membranes and cytoskeletal structures. Biophys J. 97:758-67.

San Martín, J., L. Smarandab, and T. Takahashi. 2009. Convergence of a finite element/ALE method for the Stokes equations in a domain depending on time. J. Comput. Appl. Math. 230: 521-545.

